# Phosphoproteomics data-driven signalling network inference: does it work?

**DOI:** 10.1101/2022.09.07.506895

**Authors:** Lourdes O. Sriraja, Adriano Werhli, Evangelia Petsalaki

## Abstract

The advent in high throughput global phosphoproteome profiling has led to wide phosphosite coverage and therefore the need to predict kinase substrate associations from these datasets. However, for multiple substrates, the regulatory kinase is unknown due to biased and incomplete interactome databases. In this study we compare the performance of six pairwise measures to predict kinase substrate associations using a purely data driven approach on publicly available dynamic time resolved and perturbation phosphoproteome data using mass spectrometry profiling. First, we validated the performance of these measures using as a reference both a literature-based phosphosite-specific protein interaction network and a predicted kinase substrate (KS) interactions set. The overall performance in predicting kinase-substrate associations using pairwise measures across both database-derived and predicted interactomes was poor. To expand into the wider interactome space, the performance of these measures was evaluated against a network compiled from pairs of substrates regulated by the same kinase (substrate-substrate associations). Similar to the kinase substrate predictions, a purely statistical approach to predict substrate-substrate associations was also poor. However, the addition of a sequence similarity filter for substrate-substrate associations led to a boost in performance and to the inference of statistically significant substrate-substrate associations. Our findings imply that the use of a filter to reduce the search space, such as a sequence similarity filter, can be used prior to the application of network inference methods to reduce noise and boost the signal. We also find that the current gold standard for reference sets is not adequate for evaluation as it is limited and context-agnostic. Therefore, there is a need for additional evaluation methods that have increased coverage and take into consideration the context-specific nature of kinase substrate associations.

## Introduction

Signalling networks orchestrate the cell’s responses to changes in their environment via stimuli-induced post translational modifications (PTMs) and are often deregulated in disease, including cancer, diabetes and other disorders^1^. Unravelling the connectivity of cell signalling networks can enable the understanding of the underlying mechanisms at play.

Protein phosphorylation is the best studied PTM regulating cell signalling networks and is mediated by protein kinases that transfer the ***γ****-phosphate* from ATP to Ser, Thr or Tyr residue sites on the substrate. Over 500 kinases have been identified in human cells^2^, many of which are able to phosphorylate multiple proteins and sites within the same protein.

Current knowledge about downstream targets of kinases follows a Pareto optimal power law in which 20% of the kinases accounts for 87% of substrates in the compiled interactome databases^3^. These databases are geared towards well-studied kinases resulting in an inherent bias in which certain kinases are over-represented in pathway annotations^4^. In addition, there is limited coverage of kinase substrate binding due to the difficulty in capturing interactions that are transient in nature by affinity purification methods or Y2H assays^5^. Finally, kinase substrate interactions depend on the cell state, type, mutation background and environmental context. As such, even if all putative kinase-substrates were characterised, it is impossible to systematically characterise signalling networks across all potential contexts. There is thus a need for data-driven kinase-substrate regulatory network inference from unbiased context-specific whole-cell data, such as mass spectrometry-based global phosphoproteomics data.

To explore the wider kinome space, several kinase substrate prediction methods have been developed based on amino acid position specific scoring matrices (PSSMs) of annotated substrates^6^, neural networks that combine probabilistic network from the STRING database (NetworkIN)^7^ or the use of Bayesian additive regression trees that incorporate kinase substrate edge prediction with the kinase mode of regulation (activating or inhibiting) on the phosphosite^8^. More recently prediction methods have evolved to incorporate a wider kinome space using knowledge graph-based embeddings of kinase consensus sites across individual kinases, kinase groups and families^9^. This supervised learning approach utilises predictive models that generate probabilistic scores for each prediction. These approaches rely on prior knowledge and therefore assessing the performance of these methods poses a challenge as vast regions of the signalling network remain unexplored.

High throughput phosphoproteome profiling via mass spectrometry workflows have led to increased global coverage with the identification of phosphosites exceeding 30,000 per experiment^10^. As they depict a relatively unbiased snapshot of a cell’s signalling state, they are, in theory, an excellent starting point for deriving data-driven and context-specific signalling networks.

Indeed, several methods have been developed, largely based on approaches already used in gene regulatory network inference from transcriptomics data. These fall in 6 different categories: pairwise association measures^11,12^, linear regression^13^, ordinary differential equations^14^, ensemble approaches^15,16^ and Bayesian networks^17^. These methods have been typically developed for specific datasets and whether they are generally suitable for phosphoproteomics data-driven network inference remains an open question. The HPN DREAM challenge benchmarked many methods for the task of inferring a causal network using dynamic RPPA (reverse phase protein array) datasets across different cell lines and perturbations (addition of inhibitor)^12^. The challenge^18^ concluded that the best performing methods generally take advantage of prior knowledge networks. However, RPPA data is limited compared to global phosphoproteomics, in that there are much fewer phosphosites that are profiled, and all lie in the ‘well studied’ space of cell signalling given that there are available antibodies for them. Therefore, the networks that are inferred based on this data are limited in their coverage of the global cell signalling networks.

In this study we explore the potential of pairwise association and statistical dependency measures to derive unbiased signalling networks from global phosphoproteomics datasets, i.e., networks that don’t depend on the availability of prior knowledge pathways. We focus on six pairwise and non-parametric model-based measures that are commonly used or form the basis for many other methods for data-driven inference of signalling networks from global phosphoproteomics datasets. These consist of Pearson, Spearman and Kendall correlation, mutual information, Gaussian graphical model (GGM) and FunChiSq (Methods)^19^. We provide a systematic comparison of the performance of these measures on both time series and perturbation global phosphoproteomics datasets to predict kinase-substrate associations and substrate-substrate associations (phosphosite substrates regulated by the same kinase) using a purely data driven approach (**Figure 1**).

**Figure 1:**
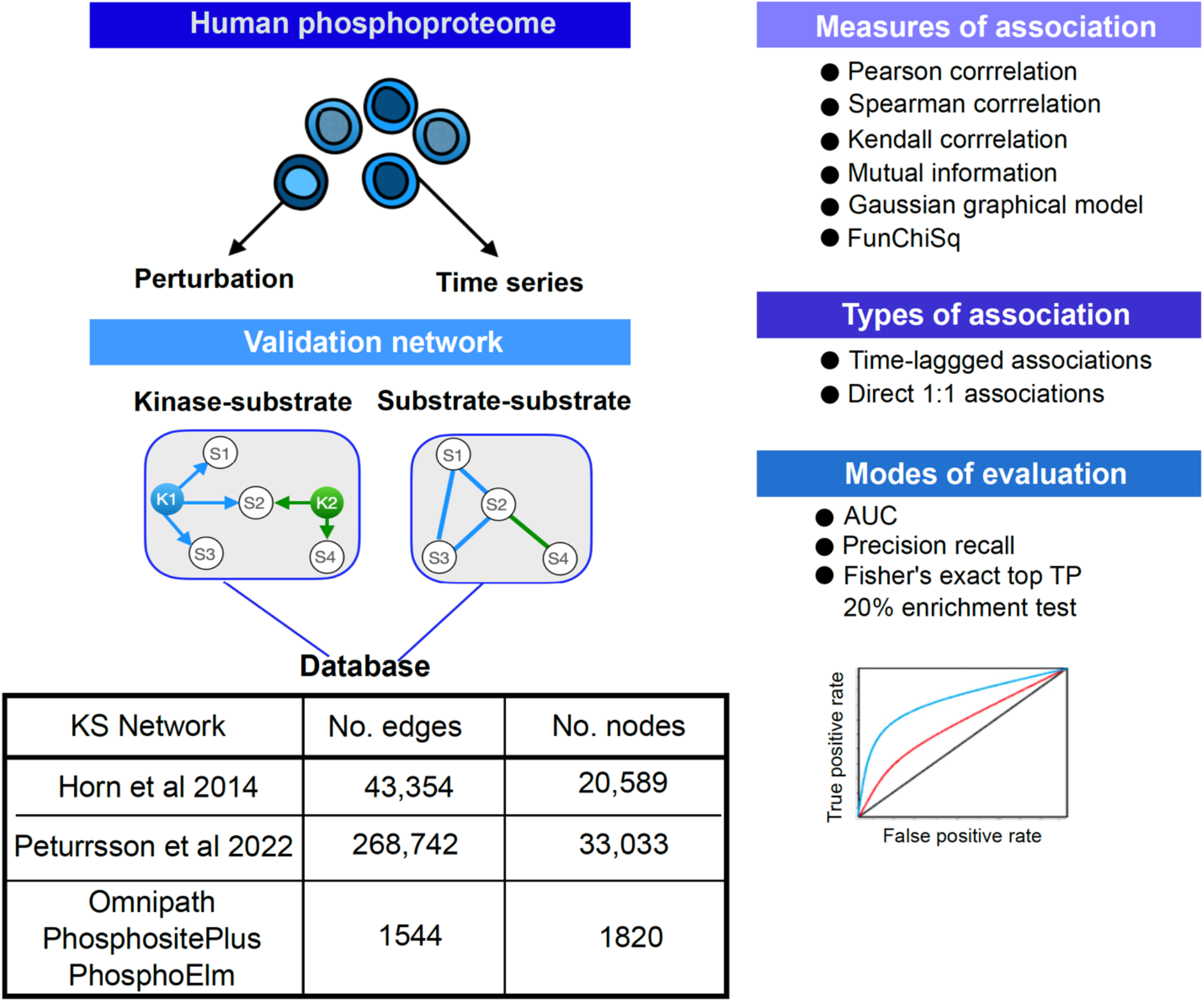
Workflow of study. We used human drug perturbation and dynamic phosphoproteome datasets and evaluated the ability of 6 metrics to identify kinase substrate associations and substrate-substrate associations. For the latter a positive edge is formed between pairs of substrates that are regulated by the same kinase. A phosphosite-specific interactome was compiled from Omnipath, PhosphositePlus and PhosphoElm. The performance of six different measures of association were evaluated on 1:1 associations for time series and drug perturbation datasets, and time lagged associations for time series datasets. This was done using AUC, precision recall and an enrichment of the number of true positives (TPs) in the top 20% of predictions.

## Results

### Simulation-based analysis suggests that number and timing of phosphoproteomics samples is critical for network inference method performance

As a first step, to set the baseline of performance for the datasets we performed a simulation study. For this, we generated synthetic data using a multivariate vector autoregressive model based on the Raf signalling pathway from Sachs et al 2005^20^, comprising 19 edges and 11 nodes (**Figure 2A**). The synthetic data was simulated with varying lengths of duration and sampled intermittently across different time intervals (t=1,2,3,4 timepoints). The performance of the six measures mentioned above were evaluated in this study to recover the underlying network.

**Figure 2:**
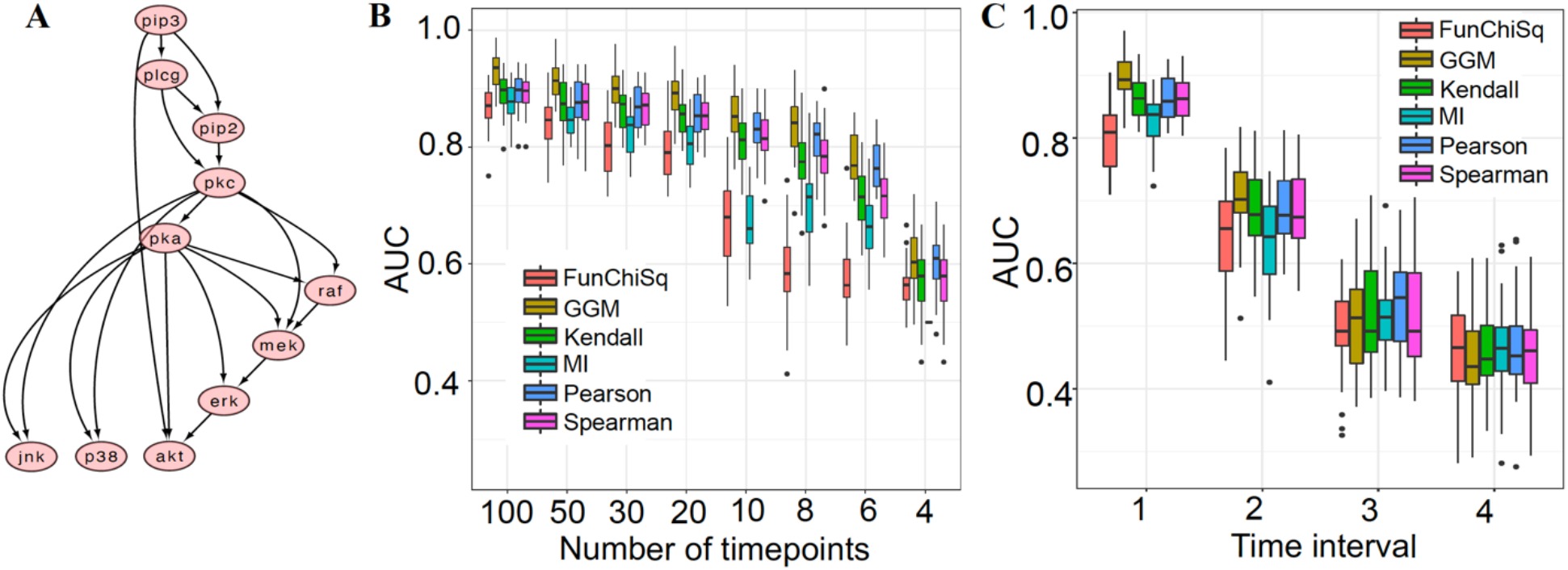
Results of kinase substrate relationship predictions on simulated data. **A**. RAF signalling pathway from Sachs et al 2005^20^ used to generate synthetic data from a multivariate autoregressive model **B**. 10,000 datasets are generated with varying lengths of duration. The box plot shows the distribution of the AUC scores for each of the predictors across varied lengths of time **C**. Synthetically generated data with time series length of 100, is sampled intermittently every 1, 2, 3, or 4 time intervals. The distribution of AUC for each of the measures is compared across the different time intervals.

Overall, we found that the AUC for all methods improves as the number of time points in the synthetic data increases (**Figure 2B**), as expected and in line with prior published analyses^21^. The curse of dimensionality (CoD), wherein a large number of proteins/genes are measured for each sample and there are more variables (nodes in the network) than timepoints, confounds the performance of any association measure. The CoD is inherent in the synthetic data at smaller time points (t=4,6,8) however the AUC performance begins to plateau after t=20, once the number of samples exceeds the number of proteins.

In addition, intermittent sampling was performed on synthetic datasets with 100 time points (**Figure 2C**). The data was either sampled every 1,2,3, or 4 time points for each of the 10,000 synthetically generated datasets with 100 time points. We observed a steep reduction in performance as the lag between the sampled time points increases. At lags of 3 and 4, the AUC value for all methods is close to or less than 0.5. More frequent sampling results in higher AUC values.

The same analysis was applied to time lagged data to generate pairwise associations between pairs of samples from t = 1, …, n-1 and t = 2, …, n where n is the length of the simulated time series datasets (**Figure S1**). Despite the underlying time dependency from the model that generates the data, all six measures were unable to recapitulate the underlying network. Increasing the length of the time series had no effect in detecting time lagged associations, similarly intermittent sampling of time points generated AUC value close to 0.5.

For the simulated datasets, GGM had the highest performance. The GGM removes spurious connections due to the conditional dependence assumption which removes the effect of all other variables to generate highly related pairwise associations. However, it should be noted that the data is simulated as a Gaussian process and therefore the GGM’s top performance in this analysis is unsurprising.

### Data-driven prediction of kinase-substrate associations

We next predicted kinase-substrate associations by applying these six measures on publicly available phosphoproteomics datasets. In total 17 datasets were compiled from 11 published papers that were acquired using label-free, TMT, SILAC or iTRAQ MS based quantification^22–32^ (Methods; **Table S1**). For the 13 time series phosphoproteome datasets, the number of sampled time points varied from four to ten. In addition, there were 4 drug perturbation datasets compiled from two studies^29,33^. Only differentially expressed phosphosites were used in this analysis (p value<0.05; log_2_ fold change > 2; Methods & **Table S1**) and evaluation results are presented only where 5 or more true positive associations were available. Three metrics were used to evaluate the performance of the six measures. Area under the curve (AUC), ratio of the area under the precision recall curve/baseline (AURPC/baseline) and Fisher’s exact odds ratio of true positive enrichment (TP) within the top 20% of the predictions.

Each of the performance metrics were initially evaluated against an interactome-derived phosphosite specific S/T kinase substrate gold network (Methods). In general, the performance was poor with no method surpassing a mean AUC of 0.6 for the S/T phosphorylation predictions. The best performing measure in detecting direct 1:1 S/T kinase substrate associations is GGM with mean AUC ∼0.6), mean log_10_(AUPRRC/baseline) =∼0.7, and mean odds ratio of TP enrichment in top 20% hits=2.5. (**Figure 3A, Figure S2A & S3A**). All methods performed slightly worse on the perturbation datasets than the time series ones, except for the FunChiSq method which performed better, though still close to random. For Y kinase substrate association predictions, the average performance is similar to that of the S/T ones, however there is a more bimodal distribution, whereby 2 datasets (Reddy et al 2016 - 10nmol, Vemulapalli et al 2021 - DMSO) consistently perform relatively well with AUCs around 0.7 whereas the rest mostly perform worse than random (**Figure 3B & S2B, S3B**). Here the GGM, Kendall, Pearson and Spearman methods performed similarly with Kendall slightly outperforming the other methods.

**Figure 3:**
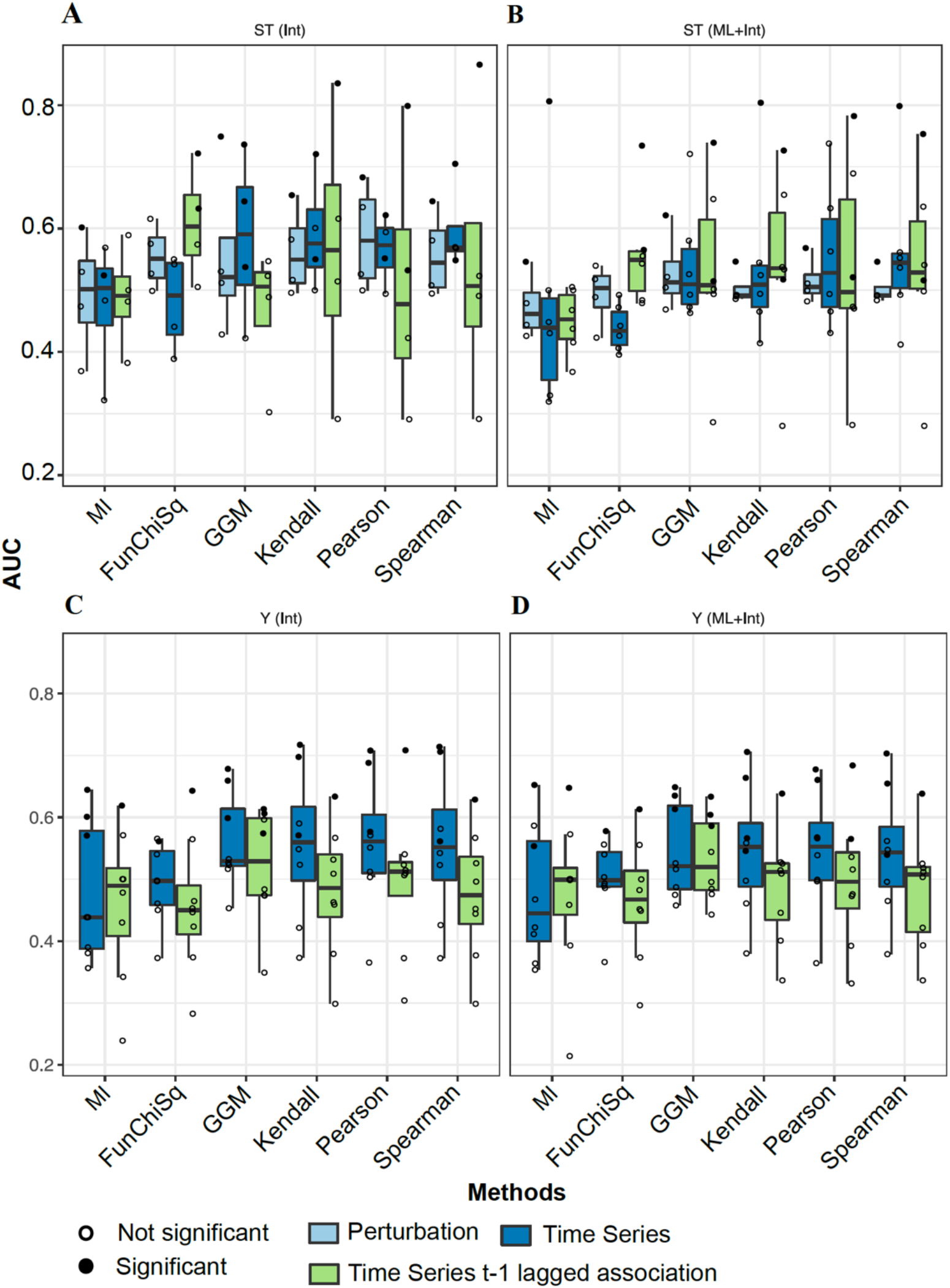
Evaluation results of kinase-substrate predictions. **A**. AUC distribution profile of S/T kinase substrate associations evaluated against database derived phosphosite specific interactions **B**. AUC distribution profile of S/T kinase substrate associations evaluated against the interactome and machine learning generated kinase substrate predictions **C**. AUC distribution profiles of Y kinase substrate associations evaluated against database derived interactome **D**. AUC distribution profile of Y kinase substrate associations evaluated against the interactome and machine learning generated kinase substrate predictions.

For time lagged S/T kinase substrate associations the FunChiSq method was the best performer (∼0.6, ∼0.6, ∼2 respectively for the evaluation metrics; **Figure 3A, Figure S2A & S3A**), while GGM performed close to random. For time lagged Y phosphorylation predictions, the performance was generally random or worse than random, except for one dataset that had a higher percentage of true positives among the possible associations (Redd et al 2016 - 10nmol) and achieved an AUC of 0.7 with the Pearson correlation metric (**Table S2**).

One of the caveats of this analysis is the low number of true positive interactions in many datasets. This meant that very few datasets were suitable for evaluation (**Table S2**). We hypothesised that increasing the number of true positives could help us better evaluate the performance of these methods by including additional datasets and putative positive points. We thus compiled kinase substrate networks from two machine learning methods that specialise in predicting kinase substrate associations at the phosphosite level (NetworKin^34^ and Selphi2.0^35^). These predictions were combined with the gold interactome to expand the potential space for evaluation, allowing us to include 2 additional datasets in the evaluation of S/T kinase substrate association predictions.

AUC distribution profile of Y kinase substrate associations evaluated against the interactome and machine learning generated kinase substrate predictions.

Using the lenient cutoff for the machine learning-inferred interactions (0.5), the 2 additional datasets (Schmutz et al 2013^31^ and Koksal et al 2018^22^) performed relatively well with the best AUCs being 0.8 with Spearman and Kendall correlation method and 0.63 with the Pearson correlation method respectively. However, the remaining datasets showed on average slightly reduced performance (**Figure 3C, S2E, S3E**), including the perturbation datasets, probably due to the relatively high level of noise expected to be included in predicted kinase-substrate associations. There was no effect on the performance of the Y kinase substrate association predictions (**Figure 3D, S2F, S3F**).

### Data-driven predicting substrates regulated by the same S/T kinase

Given the small number of true positives in kinase substrate association networks, we then asked whether predicting substrate phosphosites regulated by the same kinase would yield better performance. Pairs of substrates regulated by the same kinase were combined to form a substrate-substrate association network (**Figure 1**). The compiled gold phosphosite specific interactome for S/T phosphosites has 36,034 interactions, whilst the Y phosphosite interactome is limited to 6,641 interactions. For most datasets there were not sufficient Y phosphosites to evaluate the methods, so we decided to focus on S/T phosphosites.

We found that a purely statistical association-based prediction of substrate-substrate associations performs similarly to the kinase-substrate predictions above, with maximum mean AUCs achieved by GGM, Kendall and Spearman associations (∼0.6). The performance on the time lagged exercise or the perturbation datasets was random (**Figure 4A, S5A, S6A**).

**Figure 4:**
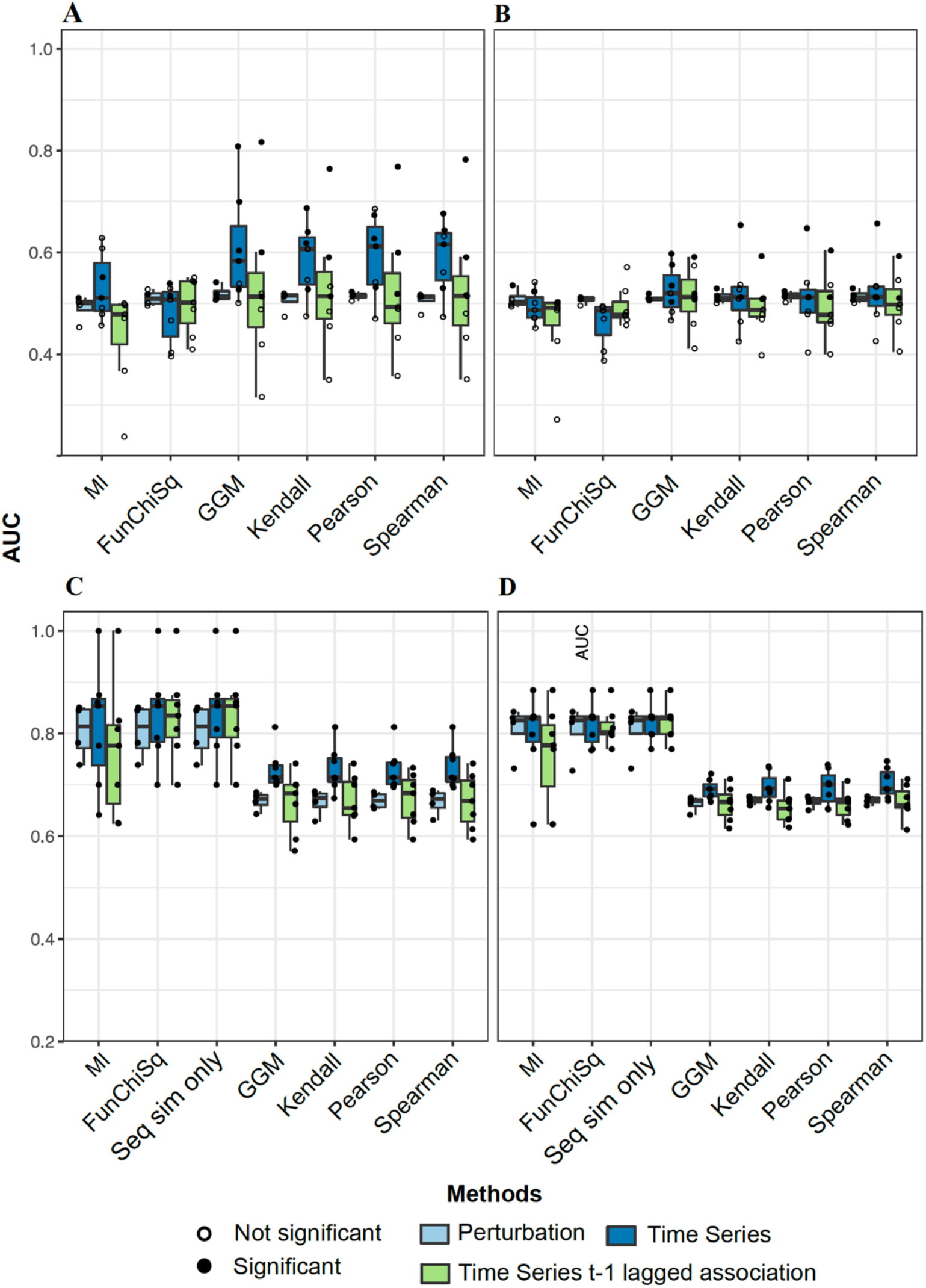
Evaluation results of substrate-substrate predictions. **A**. AUC distribution profile of substrate-substrate associations evaluated against database derived phosphosite specific interactions **B**. AUC distribution profile of substrate-substrate associations evaluated against the interactome and machine learning generated substrate-substrate predictions **C**. AUC distribution profiles of substrate-substrate associations with a sequence similarity filter evaluated against database derived interactome **D**. AUC distribution profiles of substrate-substrate associations with a sequence similarity filter evaluated against database derived interactome combined with ML derived substrate-substrate predictions

We next sought to reduce the space of false positives using our knowledge that kinases tend to recognise sites that are more similar in sequence, often matching specific motifs. By comparing the sequence similarity between known interactions and random pairs we found a small but significant increase for substrates of S/T kinases from a median of 0.142 to a median of 0.166 (**Figure S7A)**. We thus filtered all predicted association networks by setting the association score of any substrate pairs that had a sequence similarity of less than 0.166 to 0 and evaluating them as predicted negatives.

The application of sequence similarity significantly boosts the performance in generating direct substrate-substrate associations (AUCs ∼0.7-0.85), with all datasets becoming statistically significant based on the permutation analysis across all three-evaluation metrics (**Figure 4B, S5B, S6B**). The best performance here was achieved by mutual information and the FunChiSq metrics, which had been the worst performers thus far. Some AUCs achieved greater than 0.85 (**Table S2**). In fact, using the sequence similarity filtered network without any association metric and evaluating the match to the gold interactome gave the best performance across datasets (**Figures 4C-D** and **Figures S5 & S6 B, D, F**). This brings to question the value of the association measures for identifying regulatory relationships. However, even across high, medium, or low scoring associations, the difference in sequence similarity between true positives and true negatives was the same. Despite the expectation that high sequence similarity is associated with high pairwise associations this is clouded by large numbers of false positives (**Figure S7C&D**). This means that regardless of how correlated or statistically associated two substrates are, the sequence similarity is equally likely to predict them as associated in all datasets and conditions. Given that the gold set itself is agnostic to context/conditions, the sequence similarity filter can be used to identify the starting network on which association methods can be applied to identify condition/context-specific associations, while reducing the search space and thus the number of false positives.

## Discussion

A cell’s response to cell communication and external perturbation is highly context specific and is regulated by cell signalling processes. Our current understanding of these processes is based on biased and context-agnostic annotated pathways which are not accurately representative. Network inference from global phosphoproteomics data is an attractive way to extract condition-specific and relatively unbiased signalling networks.

In this study, we performed a systematic comparison of measures commonly used as the basis of most network inference methods to detect kinase substrate relationships, and pairwise substrate associations regulated by the same kinase using dynamic and perturbation phosphoproteome datasets. The goal was not to be comprehensive but to identify advantages and disadvantages of different approaches and suggest the best strategies for using them.

To create a baseline of our expectation, we started with a simulated study on synthetic data that demonstrated the known potential pitfalls of inferring associations with time series data. In particular, the length of the time series determines the performance accuracy, and intermittent sampling across the simulated data makes it difficult to recover the connections from the underlying network especially when the sampled intervals increase. This highlights the importance of time point selection when it comes to dynamic phosphoproteome profiling. Curiously, trying to identify statistical associations using a time lagged strategy was difficult both in the simulated data and in real datasets. From a biological point of view, this might be because phosphorylation reactions occur at varied timescales and the sampled timepoints don’t always reflect these reactions. In particular, tyrosine phosphorylation occurs within seconds to minutes of cell stimulation ^36^, whilst serine/threonine phosphorylation occurs over slightly longer and more varied timescales (minutes rather than seconds).

Despite the incomplete coverage and limited ground truth to evaluate the phosphoproteomics datasets, we based the evaluation of the metrics on a gold set extracted from prior knowledge databases. Although FunChiSq was one of the best performing measures in the DREAM challenge, we found that it does not perform well in predicting associations from global phosphoproteomics datasets and is consistently the worst performing measure. Mutual information is also a poor performer for smaller time series lengths although this effect is minimised as the sample length increases in the simulated data at least. Both measures require the data to be discretized, and this is challenging for datasets that have smaller numbers of samples, due to information loss in which differences/patterns between data points can become obscured once discretized. GGM, Kendall, Pearson and Spearman correlations were typically performing similarly with GGM often performing marginally better. Throughout the study time series datasets tended to perform similarly or better than perturbation datasets, implying that collection of time series might be preferable for network inference studies.

There were instances in which the performance of certain datasets was acceptable, however overall, the performance of all metrics was poor. AUCs were in the 0.6 range and approximate log_10_ AUPR/baseline ratio of 0.7 for S/T kinase substrate predictions and only marginally higher than zero for Y kinase substrate predictions. Whilst true positive interactions were indeed associated with higher scores of associations, it was clouded by the overwhelming number of false positive hits. This is because most of these association metrics don’t necessarily reflect causal relationships, even though it is expected that if they indeed are causal, they will also have high association scores. A major consideration is also the suitability of the prior knowledge-derived gold dataset that is commonly used in the evaluation of network inference methods. We observed that the majority of phosphosites had very low numbers of true positives available for discovery which resulted in only 4-5/13 of our time series datasets being used in the evaluation. This reflects the vast dark space of the human signalling networks^3,4^, and makes evaluating connectivity using noisy phosphoproteome datasets challenging as it is impossible to know whether high scoring associations are indeed true or false positives. This was true even when we attempted to generate networks of correlated putative kinase substrates: whilst we were able to increase the number of true positives available for evaluation, the potential false positives also increased subsequently. Using machine learning based predicted kinase substrate relationships to extend our gold set resulted in the inclusion of additional datasets in the evaluation that performed relatively well, but the introduction of the noise of these predicted sets, mostly reduced the performance of datasets that already had a high coverage of true positives.

The introduction of a sequence similarity filter between associated substrates, enhanced the performance leading to AUCs in the order of 0.8 or even higher. This was only possible for S/T kinase substrates as there was no shift in the distribution of the sequence similarity of Y kinase substrates (**Figure S7A&B**), possibly due to lack of extensive motif specificity ^37^ or alternative substrate specificity strategies through e.g. SH2 domains^38^.

The FunChiSq and mutual information were top performing metrics in par with the sequence similarity filter as a sole measure. A possible hypothesis for this boost in performance could be due to the inefficiency of the metrics as sole measures in predicting substrate-substrate associations (**Figure 4A&B**), both MI and FunChiSq perhaps become non-redundant once the filter is applied. MI and FunChiSq are discretized measures of association and rely on many samples to work effectively. In order to estimate the joint distribution for MI, the underlying distribution needs to be sampled extensively especially for a ‘plug-in’ MI estimator based on the frequency of discretized samples. Insufficient number of samples leads to overestimation of MI and therefore introduces a bias in the estimate of statistical dependency between two variables^39–41^. Although the number of samples was limited for time series datasets (approximately 4-8 samples) in this study, the number of samples for perturbation data was higher on the scale of approximately 40-60 samples across the 4 datasets. However, the performance did not improve for the perturbation datasets, and the AUC distribution remained around 0.5 (**Fig 4A&B)**.

Recently, a method that performs dual data and phosphosite motif clustering showed promising results in network inference from phosphoproteomics data^42^. This corroborates the value of introducing a sequence similarity-based feature in network inference strategies and highlights the importance of reducing the noise in a dataset before applying network inference methods. In addition, efforts have recently been made to identify the substrate recognition motifs for all kinases^43,44^, which will further improve our ability to reduce noise and match kinases to the correct substrates. Additional ways to reduce noise in the dataset could include filters to include phosphosites that are predicted to be functional^45^, and introducing very strict p-value and fold-change cut-offs for the identification of differentially abundant phosphosites. Interestingly the filtered sequence similarity metric as a sole measure performed better than in combination with an association metric. This reveals an additional issue with the current ways of evaluating these methods using prior knowledge gold sets, as they are context-agnostic and represent a sum of putative relationships that have been observed in vastly different cell types, stimuli, and other conditions. Therefore, any method to identify associations in each phosphoproteomics dataset assumes that these associations exist if the relevant proteins are there, regardless of the conditions. However, this assumption is not always true and could contribute towards the apparent poor performance of the methods.

Indeed, we found that the sequence similarity filter metric was similar across all levels of association scores (**Fig S7C&D**), even though true positives tend to have higher such scores. Thus, it seems suitable to combine association metrics that provide the data-specific context, with strategies to reduce noise such as such functionality, sequence similarity or motif-based filtering to extract data driven and context specific kinase-substrate networks from phosphoproteomics datasets.

In conclusion, network inference approaches based on the six association metrics evaluated in this study doesn’t perform well when used as a sole measure when evaluated against prior knowledge-derived gold sets of kinase substrate relationships. However, methods that can a priori reduce the noise and boost the signal, such as a sequence similarity filter boosts the performance substantially.

A major issue when evaluating these methods is the sparsity of the gold set. There are several efforts underway to shed light on the dark space of kinase substrate regulatory networks^3,8,35,43,44,46,47^, which should largely mitigate the limitation of low coverage in prior knowledge-derived gold sets to evaluate network inference methods. However, these gold sets will remain context agnostic, making it difficult to identify the true performance of network inference methods. There is thus a need for approaches that combine performance metrics on more comprehensive gold sets, with evaluation based on orthogonal datasets.

## Code and data

All code required for reproducing this work is freely available at https://sandbox.zenodo.org/record/1100183#.YxZ229LMJhE

## Funding

AW was funded by the CABANA secondment program, which is funded by Global Challenges Research Fund - part of the UK AID budget. LS was funded by the EIPOD3 program which is supported by the Marie Skłodowska-Curie Cofund programme. The project was also supported by core funding provided by EMBL-EBI.

## Credit authorship contribution statement

Conceptualization: EP, LS, AW, Methodology: LS, EP, AW, Software: LS, AW, Validation: LS, AW, Formal analysis: LS, AW, Data Curation: LS, Writing - original draft: LS, AW, Writing - review and editing: LS, EP, Visualization: LS, Supervision: EP, Project administration: EP, Funding acquisition: LS, AW, EP.

## Declaration of Competing Interest

The authors declare that they have no known competing financial interests or personal relationships that could have appeared to influence the work reported in this paper.

## Materials & Methods

### Selection of datasets and methods to evaluate

We selected the kinase substrate association methods to evaluate in this study as follows: The most commonly utilised pairwise expression measure omic-wide is Pearson correlation, which is used to generate co-expression networks of functionally relevant proteins/genes^12,48^. Rank based Spearman and Kendall correlation were found to be the best performing measures in identifying co-expressed genes from the same pathway and were therefore selected as a comparative measure in this analysis^49^. Mutual information-based methods aim to capture nonlinear dependence and have been shown to perform well in recapitulating gene networks^50,51^ and frequently used to reconstruct gene regulatory networks. FunChiSq is a more recently derived directional statistical test of association based on the asymptotic null chi-squared distribution^52^. As part of the HPN-DREAM challenge^18^, FunChiSq was highly ranked in detecting protein associations from RPPA experimental data as well as the top performer in predicting associations from simulated model data. Finally, the partial correlation based Gaussian graphical model (GGM) is able to detect the association of two genes/proteins given this pair are conditionally dependent on the remaining genes/proteins in the data^53^.

All phosphoproteome datasets were selected based on being derived from human cell lines to ensure wider interactome coverage, and sufficient number of samples for evaluation of associations. For dynamic time series datasets, the data needed to be sampled across 4 or more time points, the perturbation datasets used in this analysis had sufficient samples for evaluation. Both the time series and perturbation datasets were filtered for log2 fold change in expression in relation to control or t=0.

### Data processing

Individual datasets (phosphoproteome and proteome) were processed differently depending on the MS profiling approach used. TMT datasets were column median and row mean normalized to minimise batch effect variations across the replicates. SILAC datasets were normalised to control samples as ratios. Label free datasets were available pre-processed. In addition, some of the published datasets did not have replicates whilst other datasets only included the average intensity across the replicates. The detailed explanation on data processing specific to each dataset is provided in **Table S1**.

Post-processing batch effects for data with biological or technical replicates were tested using the procedures outlined in Nusinow et al 2020^54^ (PCA, Hierarchical clustering and linear models). Datasets that exhibited batch effects post-processing were not included in this analysis. Dynamic phosphoproteome datasets with time points that exceeded 1 hour were normalized to the corresponding proteome as phosphoproteome to proteome ratios.

The spline based linear mixed model (LMMS package in R) package in R was used to identify differentially expressed phosphosites across time series datasets. Only datasets from human cell lines or patients were analysed in this study.

Differential expression analysis was performed using the *lmmsDE* function in R for datasets that include replicates^55^, where a linear mixed model spline is fit to the data and compared to a null model that fits to the intercept. The function returns Benjamini-Hochberg corrected p-values and an adjusted p-value threshold of 0.05 was used to filter for significant profiles. The phosphosites across replicates were averaged and an additional fold change (FC) filter to each phosphosite response is applied in which phosphosite log2 ratios to t=0 has at least one time point with |log2 FC| > 2. Published datasets that only included the average phosphosite signal or datasets with no replicates were filtered with |log2 FC| > 2.

Two perturbation datasets were also used as a comparison. The published perturbation datasets from Wilkes et al 2015 and Hijazi et al 2020^29,33^ were already pre-processed as log2 ratios with respect to controls and each phosphosite in each condition has an associated p-value. The perturbation datasets were filtered based on the following: each phosphosites that had 20% or more p-values<0.05 and 2 or more phosphosites with |log2 FC| > 2.

### Association measures

Pearson correlation coefficients are computed from continuous data and capture linear associations. Given two time series *x=*(*x*_1_,…,*x*_*k*_) and *y=*(*y*_1_,…,*y*_*k*_) each of length k, the Pearson correlation coefficient between these variables is:

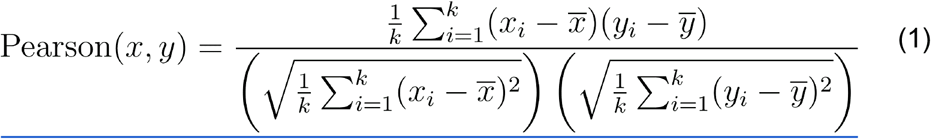

where 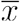 and 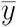 are the empirical means of *x* and *y* respectively.

Kendall’s Tau association is a non-parametric rank-based measure between two variables that is able to capture nonlinear dependencies. Given *x*_*i*_ and *y*_*i*_ are unique, any pair of observations (*x*_*i*_,*y*_*i*_) and (*x*_*i*_,*y*_*i*_) are classified as concordant if the ranks of both elements agree, i.e., rg(*x*_*i*_)= rg(*x*_*j*_) and rg(*y*_*i*_)= rg(*y*_*j*_) (rg(.) denotes the rank), and are discordant if they do not agree. The number of concordant and discordant pairs is then employed to calculate the Kendall association score:

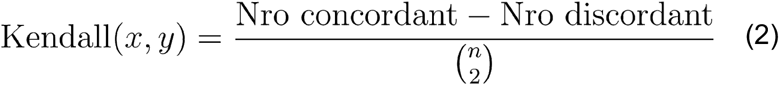

Spearman correlation coefficient is a non-parametric rank correlation score between two variables. Spearman correlation is obtained by calculating the Pearson correlations between the rank of the variables:

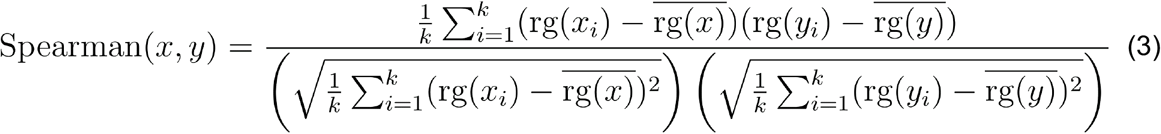

Mutual information is a non-parametric measure of linear or non-linear association derived from information theory. Both x and y variables are discretized, where *p*(*x=i, y=i*) is the joint probability of *x* and *y*. Given that variables *x* and *y* are discretized in *r* levels the MI between these variables can be defined as:

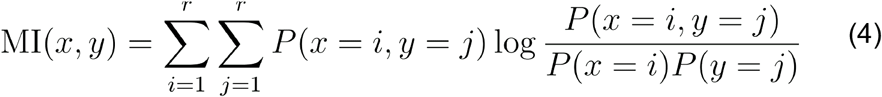

The functional Chi-squared test is a more recent measure of non-parametric causal dependency based on the normalized Chi-square test^19,52^. Each time series is first discretized by 1d k-means clustering using the Ckmeans.1d.dp function from the Ckmeans.1d.dp package in R^56^. A contingency table is generated and used as an input into the fun.chisq.test function that generates a normalized chi square statistic as a measure of pairwise dependence. This is the only non-symmetric measure used in this study that is able to detect a directional association as it is based on the asymptotic null chi-square distribution.

In the context of relevance networks, a *n*×*n* matrix represents all the interactions between *n* nodes, and the entries are computed as either Person (*m*_*i*_,*m*_*j*_), Kendall(*m*_*i*_,*m*_*j*_),Spearman(*m*_*i*_,*m*_*j*_) and MI(*m*_*i*_,*m*_*j*_).

All the above measures are pairwise associations, however a Graphical Gaussian model (<u>GGMs</u>) is a multivariate approach with the assumption that the data is normally distributed. This forms the basis of undirected probabilistic graphical models with conditional dependencies across all the variables. GGMs are based on a (stable) estimation of the covariance matrix under the assumption that the data derives from a multivariate distribution. The element *C*_*ik*_ of the covariance matrix C is related to the correlation coefficient between nodes *X*_*i*_ and *X*_*k*_. A high correlation coefficient between two nodes may indicate a direct interaction, an indirect interaction, or a joint regulation by a common (possibly unknown) factor. However, only the direct interactions are used to generate associations. The strengths of these direct interactions are measured by the partial correlation coefficient *ρ*_*ik*_, which describes the correlation between nodes *X*_*i*_ and *X*_*k*_ conditional on all the other nodes in the network.

Based on the theory of normal distributions, the matrix *ρ* of partial correlation coefficients *ρ*_*ik*_ is the inverse of the covariance matrix C C^−1^, (with elements 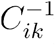):

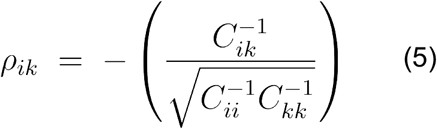

The covariance matrix is estimated using a regularized shrinkage approach outlined in Shaefer et al 2004^53^.

Direct (1:1) and time lagged associations (one unit of time) were generated for time series data, and only direct (1:1) associations were generated for the perturbation datasets.

### Simulation analysis

In order to assess the performance of the various measures used in this study on a controlled dataset and to evaluate the effect of the number and timing of samples collected, a multivariate model with fixed coefficients was used to generate dynamic synthetic data. The model assumes that the data is sampled from a white noise Gaussian process. An example of a set of equations used to describe a simple three node network for nodes A, B and C (X_A_ → X_B_, X_A_ → X_C,_ X_B_ → X_C_) is shown below:

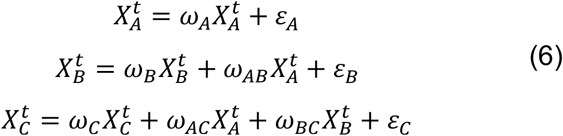

The strength of an interaction is defined by *ω* and the added noise to the simulated data is defined by *ε*. The data simulated in this paper is based on the Raf signalling pathway from Sachs et al 2005^20^, which has 11 nodes and 19 edges (**Figure 2A**). The general linear Gaussian distribution function used to generate the simulated data is based on the following equation:

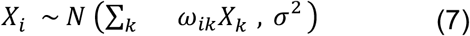

in which *N*(.) denotes the normal distribution and the sum extends over all parents of node *X*_*i*_, *X*_*k*_ represents the interacting node, with *ω*_*ik*_ representing the strength of an interaction between the interacting nodes whilst *σ*^2^ represents noise that has been added to the data.

### Evaluation of performance

The first adjacency matrix is phosphosite specific at the substrate level and consists of kinase uniprot id’s and substrate Uniprot id’s with the phosphorylation site and is compiled from PhosphositePlus^57^, Omnipath^58^ and PhosphoElm^59^.

The adjacency matrix was filtered for kinase-substrate interactions only, in which an interaction in the adjacency matrix has an entry of 1 or 0 for tyrosine kinase phosphosite rows that intersect with tyrosine phosphosites only, the remaining entries in the row are set to NA. Similarly, serine/threonine kinases have an entry of 1 (if this interaction exists in the interactome) or 0 (true negative) at the intersection to serine/threonine phosphosite substrates. Non-kinase - non-kinase interactions were assigned NAs in the adjacency matrix and were not included in the final evaluation. Performance of each association method was evaluated using receiver operating curve (ROC) precision recall (AUROC) and enrichment of true positives in top 20% predictions using Fisher’s exact test.

To overcome the limitations of understudied regions of the interactome, a kinase substrate network was compiled from two machine learning algorithms using the phosphosites across all the datasets used in this study. NetworKin from KinomeXplorer^34^ uses a naive bayes approach that combines consensus sequence motifs of kinases and their binding domains based on a model derived probabilistic score ^7^ with a string database protein proximity score to generate a final score that represents how likely the phosphorylation interaction would occur.

These predictions were combined with kinase substrate predictions from SELPHI2, an algorithm with increased coverage of kinases compared to NetworKin^35^. This method utilises a random forest model that uses position weight matrices for the kinases compiled from PhosphositePlus^57^, coexpression from human atlas and GTEX, co-phosphorylation from CPTAC breast cancer samples and kinase inhibition profiles across different cell lines^33^. In addition, this method uses features from the phosphosite functional score from Ochoa et al 2019^45^ to generate kinase substrate predictions with a probability score of how likely the kinase will phosphorylate the substrate. There were two probabilistic cut-offs used to generate two networks for each datasets at probability cut-off >0.5 and >0.8 (high confidence network).

Since phosphosites regulated by the same kinase have been shown to cluster together^60^, substrate-substrate interactions were generated to account for associations between pairs of substrates regulated by the same kinase. These associations were compiled using the same interactome databases and machine learning derived predictions as described above.

### Sequence Similarity calculations

Phosphosites regulated by the same kinase have been shown to cluster together in time series phosphoproteome datasets^60^. This could be due to multiple substrates in close proximity to the kinase, but also due to the specificity of certain kinases to its substrate phosphosites. In order to test kinase specificity to substrates, substrate phosphosites compiled from the interactome were divided into two groups: Y phosphopeptides and S/T phosphopeptides. TP interactions are generated by pairwise combinations of the substrates regulated by the same kinase, and TN interactions are generated from substrate phosphosite pairs that are not known to be regulated by the same kinase (**Fig S7A&B**).

The sequence similarity between the substrate phosphopeptides across both sets (TPs and TNs) are calculated using the parSeqSim function from the protr package in R^61^, based on the BLOSUM62 substitution matrix using a local alignment of pairs of sequence peptides each of length 15 amino acids long. The distribution of sequence similarity between pairs of substrate phosphopeptides was compared across TP and TN interactions. For Y phosphopeptides there was an overlap between the distributions of sequence similarity, however for S/T phosphopeptide pairs there was a shift to the right in the distribution of sequence similarity for TP interactions. Based on the median of the TP distribution, a 0.166 cut-off was applied to filter pairs of substrate substrate S/T associations based on their sequence similarity. The values that are filtered out are assigned a value of 0 and evaluated as a TN.

### Significance Testing

To assess the performance of different evaluation methods compared to a null hypothesis, an edge list that consisted of row column indices, an interaction column (1/0) and a measure of association between the indexed phosphosites was generated. The association measure column was randomly sampled without replacement 10,000 times to generate 10,000 randomly sampled networks. The performance of these randomly sampled networks was evaluated using the same procedures as the original datasets. A distribution of AUC values was generated for each of these randomly sampled edge lists that correspond to a dataset, measure and adjacency matrix (for e.g Biggelaar et al 2014 - Pearson correlation - S/T kinase substrate adjacency matrix). A p-value was generated using the following equation:

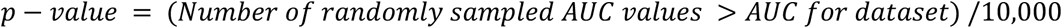

## Notes

https://sandbox.zenodo.org/record/1100183#.YxZ229LMJhE

### Competing Interest Statement

The authors have declared no competing interest.

## References

1. Chen, L., Liu, S. & Tao, Y. Regulating tumor suppressor genes: post-translational modifications. Sig Transduct Target Ther 5, 1–25 (2020).

2. Manning, G., Whyte, D. B., Martinez, R., Hunter, T. & Sudarsanam, S. The Protein Kinase Complement of the Human Genome. Science 298, 1912–1934 (2002).

3. Needham, E. J., Parker, B. L., Burykin, T., James, D. E. & Humphrey, S. J. Illuminating the dark phosphoproteome. Sci. Signal. 12, eaau8645 (2019).

4. Invergo, B. M. & Beltrao, P. Reconstructing phosphorylation signalling networks from quantitative phosphoproteomic data. Essays Biochem 62, 525–534 (2018).

5. Davey, N. E. et al. Discovery of short linear motif-mediated interactions through phage display of intrinsically disordered regions of the human proteome. The FEBS Journal 284, 485–498 (2017).

6. Miller, M. L. et al. Linear Motif Atlas for Phosphorylation-Dependent Signaling. Sci Signal 1, ra2 (2008).

7. Linding, R. et al. Systematic Discovery of In Vivo Phosphorylation Networks. Cell 129, 1415–1426 (2007).

8. Invergo, B. M. et al. Prediction of Signed Protein Kinase Regulatory Circuits. Cell Syst 10, 384-396.e9 (2020).

9. Nováček, V. et al. Accurate prediction of kinase-substrate networks using knowledge graphs. PLOS Computational Biology 16, e1007578 (2020).

10. Skowronek, P. et al. Rapid and in-depth coverage of the (phospho-)proteome with deep libraries and optimal window design for dia-PASEF. 2022.05.31.494163 Preprint at https://doi.org/10.1101/2022.05.31.494163 (2022).

11. Liu, Z.-P. Quantifying Gene Regulatory Relationships with Association Measures: A Comparative Study. Front Genet 8, 96 (2017).

12. Langfelder, P. & Horvath, S. WGCNA: an R package for weighted correlation network analysis. BMC Bioinformatics 9, 559 (2008).

13. Dong, Z., Song, T. & Yuan, C. Inference of Gene Regulatory Networks from Genetic Perturbations with Linear Regression Model. PLOS ONE 8, e83263 (2013).

14. Ma, B., Fang, M. & Jiao, X. Inference of gene regulatory networks based on nonlinear ordinary differential equations. Bioinformatics 36, 4885–4893 (2020).

15. Ruyssinck, J. et al. NIMEFI: Gene Regulatory Network Inference using Multiple Ensemble Feature Importance Algorithms. PLOS ONE 9, e92709 (2014).

16. Huynh-Thu, V. A., Irrthum, A., Wehenkel, L. & Geurts, P. Inferring Regulatory Networks from Expression Data Using Tree-Based Methods. PLOS ONE 5, e12776 (2010).

17. Xing, L. et al. An improved Bayesian network method for reconstructing gene regulatory network based on candidate auto selection. BMC Genomics 18, 844 (2017).

18. Hill, S. M. et al. Inferring causal molecular networks: empirical assessment through a community-based effort. Nature Methods 13, 310–318 (2016).

19. Zhong, H. & Song, M. A Fast Exact Functional Test for Directional Association and Cancer Biology Applications. IEEE/ACM Transactions on Computational Biology and Bioinformatics 16, 818–826 (2019).

20. Sachs, K., Perez, O., Pe’er, D., Lauffenburger, D. A. & Nolan, G. P. Causal Protein-Signaling Networks Derived from Multiparameter Single-Cell Data. Science 308, 523–529 (2005).

21. Zou, C. & Feng, J. Granger causality vs. dynamic Bayesian network inference: a comparative study. BMC Bioinformatics 10, 122 (2009).

22. Köksal, A. S. et al. Synthesizing Signaling Pathways from Temporal Phosphoproteomic Data. Cell Reports 24, 3607–3618 (2018).

23. Vemulapalli, V. et al. Time-resolved phosphoproteomics reveals scaffolding and catalysis-responsive patterns of SHP2-dependent signaling. eLife 10, e64251 (2021).

24. Ji, Q., Ding, Y. & Salomon, A. R. SRC Homology 2 Domain-containing Leukocyte Phosphoprotein of 76 kDa (SLP-76) N-terminal Tyrosine Residues Regulate a Dynamic Signaling Equilibrium Involving Feedback of Proximal T-cell Receptor (TCR) Signaling. Mol Cell Proteomics 14, 30–40 (2015).

25. Song, Y. et al. A dynamic view of the proteomic landscape during differentiation of ReNcell VM cells, an immortalized human neural progenitor line. Sci Data 6, 190016 (2019).

26. van den Biggelaar, M. et al. Quantitative phosphoproteomics unveils temporal dynamics of thrombin signaling in human endothelial cells. Blood 123, e22–e36 (2014).

27. de Graaf, E. L. et al. Signal Transduction Reaction Monitoring Deciphers Site-Specific PI3K-mTOR/MAPK Pathway Dynamics in Oncogene-Induced Senescence. J. Proteome Res. 14, 2906–2914 (2015).

28. Reddy, R. J. et al. Early signaling dynamics of the epidermal growth factor receptor. Proc Natl Acad Sci U S A 113, 3114–3119 (2016).

29. Wilkes, E. H., Terfve, C., Gribben, J. G., Saez-Rodriguez, J. & Cutillas, P. R. Empirical inference of circuitry and plasticity in a kinase signaling network. PNAS 112, 7719–7724 (2015).

30. Hijazi, M., Smith, R., Rajeeve, V., Bessant, C. & Cutillas, P. R. Reconstructing kinase network topologies from phosphoproteomics data reveals cancer-associated rewiring. Nature Biotechnology 38, 493–502 (2020).

31. Schmutz, C. et al. Systems-Level Overview of Host Protein Phosphorylation During Shigella flexneri Infection Revealed by Phosphoproteomics *. Molecular & Cellular Proteomics 12, 2952–2968 (2013).

32. Chiang, C.-K. et al. Quantitative phosphoproteomics reveals involvement of multiple signaling pathways in early phagocytosis by the retinal pigmented epithelium. Journal of Biological Chemistry 292, 19826–19839 (2017).

33. Hijazi, M., Smith, R., Rajeeve, V., Bessant, C. & Cutillas, P. R. Reconstructing kinase network topologies from phosphoproteomics data reveals cancer-associated rewiring. Nat Biotechnol (2020) doi:10.1038/s41587-019-0391-9.

34. Horn, H. et al. KinomeXplorer: an integrated platform for kinome biology studies. Nature Methods 11, 603–604 (2014).

35. Petursson, B. & Petsalaki, E. Data-driven extraction of human kinase-substrate relationships from omics datasets. 2022.01.15.476449 Preprint at https://doi.org/10.1101/2022.01.15.476449 (2022).

36. June, C. H., Fletcher, M. C., Ledbetter, J. A. & Samelson, L. E. Increases in tyrosine phosphorylation are detectable before phospholipase C activation after T cell receptor stimulation. J Immunol 144, 1591–1599 (1990).

37. Corwin, T. et al. Defining human tyrosine kinase phosphorylation networks using yeast as an in vivo model substrate. Cell Syst 5, 128-139.e4 (2017).

38. Miller, W. T. Determinants of Substrate Recognition in Nonreceptor Tyrosine Kinases. Acc Chem Res 36, 393–400 (2003).

39. Panzeri, S. & Treves, A. Analytical estimates of limited sampling biases in different information measures. Network 7, 87–107 (1996).

40. Samengo, I. Estimating probabilities from experimental frequencies. Phys Rev E Stat Nonlin Soft Matter Phys 65, 046124 (2002).

41. Hernández, D. G. & Samengo, I. Estimating the Mutual Information between Two Discrete, Asymmetric Variables with Limited Samples. Entropy (Basel) 21, 623 (2019).

42. Creixell, M. & Meyer, A. S. Dual data and motif clustering improves the modeling and interpretation of phosphoproteomic data. Cell Reports Methods 2, 100167 (2022).

43. Johnson, J. L. et al. A global atlas of substrate specificities for the human serine/threonine kinome. 2022.05.22.492882 Preprint at https://doi.org/10.1101/2022.05.22.492882 (2022).

44. Li, A., Voleti, R., Lee, M., Gagoski, D. & Shah, N. H. High-throughput profiling of sequence recognition by tyrosine kinases and SH2 domains using bacterial peptide display. 2022.08.01.502334 Preprint at https://doi.org/10.1101/2022.08.01.502334 (2022).

45. Ochoa, D. et al. The functional landscape of the human phosphoproteome. Nat Biotechnol 1–9 (2019) doi:10.1038/s41587-019-0344-3.

46. Berginski, M. E. et al. The Dark Kinase Knowledgebase: an online compendium of knowledge and experimental results of understudied kinases. Nucleic Acids Research 49, D529–D535 (2021).

47. Moret, N. et al. A resource for exploring the understudied human kinome for research and therapeutic opportunities. 2020.04.02.022277 Preprint at https://doi.org/10.1101/2020.04.02.022277 (2021).

48. Mantini, G. et al. Co-expression analysis of pancreatic cancer proteome reveals biology and prognostic biomarkers. Cell Oncol. 43, 1147–1159 (2020).

49. Kumari, S. et al. Evaluation of Gene Association Methods for Coexpression Network Construction and Biological Knowledge Discovery. PLoS One 7, e50411 (2012).

50. Margolin, A. A. et al. ARACNE: An Algorithm for the Reconstruction of Gene Regulatory Networks in a Mammalian Cellular Context. BMC Bioinformatics 7, S7 (2006).

51. Lachmann, A., Giorgi, F. M., Lopez, G. & Califano, A. ARACNe-AP: gene network reverse engineering through adaptive partitioning inference of mutual information. Bioinformatics 32, 2233–2235 (2016).

52. Zhang, Y. & Song, M. Deciphering Interactions in Causal Networks without Parametric Assumptions. 1311.2707 [q-bio] (2013).

53. Schafer, J. & Strimmer, K. An empirical Bayes approach to inferring large-scale gene association networks. Bioinformatics 21, 754–764 (2005).

54. Nusinow, D. P. & Gygi, S. P. A Guide to the Quantitative Proteomic Profiles of the Cancer Cell Line Encyclopedia. http://biorxiv.org/lookup/doi/10.1101/2020.02.03.932384 (2020) doi:10.1101/2020.02.03.932384.

55. Straube, J., Gorse, A.-D., Huang, B. E. & Lê Cao, K.-A. A Linear Mixed Model Spline Framework for Analysing Time Course ‘Omics’ Data. PLoS One 10, (2015).

56. Wang, H. & Song, M. Ckmeans.1d.dp: Optimal k-means Clustering in One Dimension by Dynamic Programming. R J 3, 29–33 (2011).

57. Hornbeck, P. V. et al. 15 years of PhosphoSitePlus®: integrating post-translationally modified sites, disease variants and isoforms. Nucleic Acids Res 47, D433–D441 (2019).

58. Türei, D., Korcsmáros, T. & Saez-Rodriguez, J. OmniPath: guidelines and gateway for literature-curated signaling pathway resources. Nature Methods 13, 966 (2016).

59. Dinkel, H. et al. Phospho.ELM: a database of phosphorylation sites—update 2011. Nucleic Acids Res 39, D261–D267 (2011).

60. Domanova, W. et al. Unraveling Kinase Activation Dynamics Using Kinase-Substrate Relationships from Temporal Large-Scale Phosphoproteomics Studies. PLOS ONE 11, e0157763 (2016).

61. Xiao, N., Cao, D.-S., Zhu, M.-F. & Xu, Q.-S. protr/ProtrWeb: R package and web server for generating various numerical representation schemes of protein sequences. Bioinformatics 31, 1857–1859 (2015).

