## Supplementary Figures 1-10 and Table S1 for "Phosphoproteomics data-driven signalling network inference: does it work?"

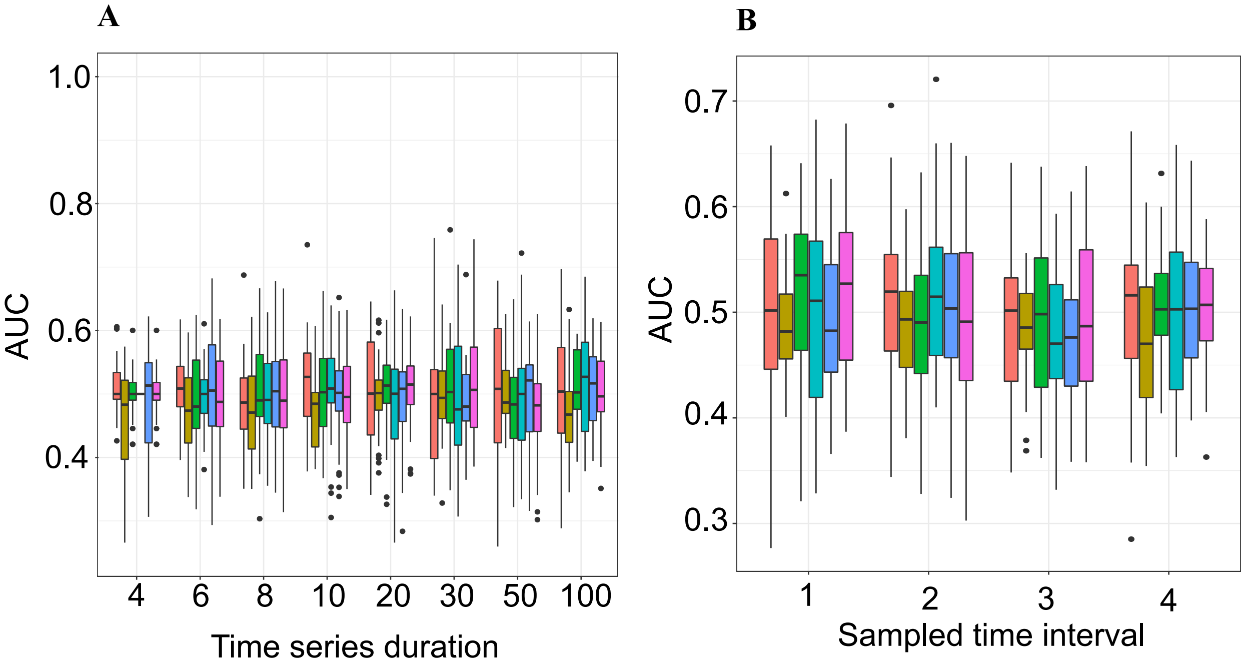


**Figure S1. Time lagged associations from the synthetic data. A.** AUC distribution when data was sampled using varying time duration as displayed in the x axis **B.** AUC distributions when we performed intermittent sampling every 1,2,3 or 4 timepoints from data of length 100 timepoints

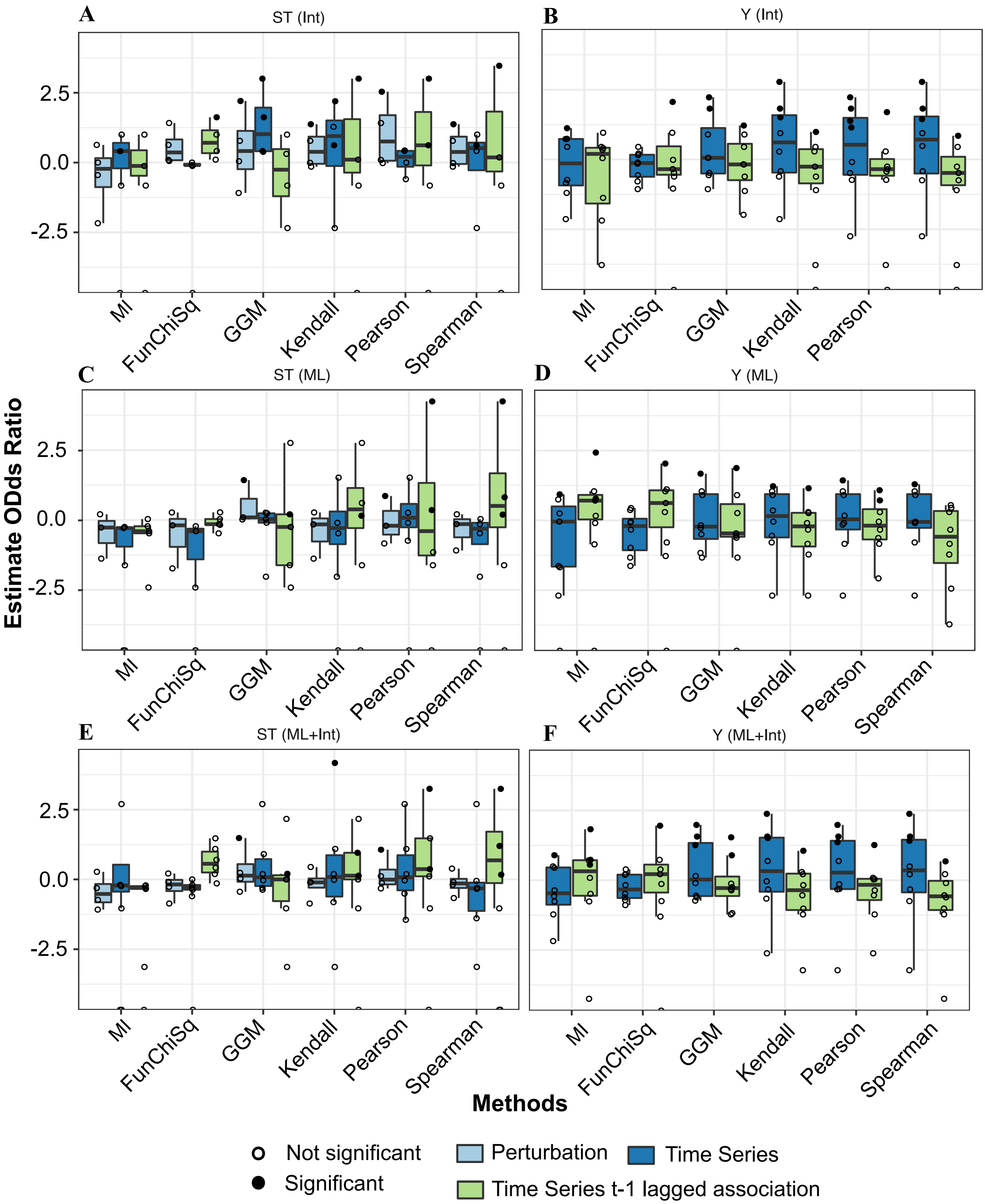


**Figure S2. Distribution plots of log10 AUPRC/Baseline ratio for interactome derived association**. **A.** S/T kinase substrate associations **B.** Y kinase substrate associations **C.** Machine learning derived S/T kinase substrate associations **D.** Machine learning derived Y kinase substrate associations. **E.** Combined interactome and machine learning derived S/T kinase substrate associations **F.** Combined interactome and machine learning derived Y kinase substrate associations. The black filled in points on the boxplots indicates statistical significance from the permutation analysis with p-value < 0.05.

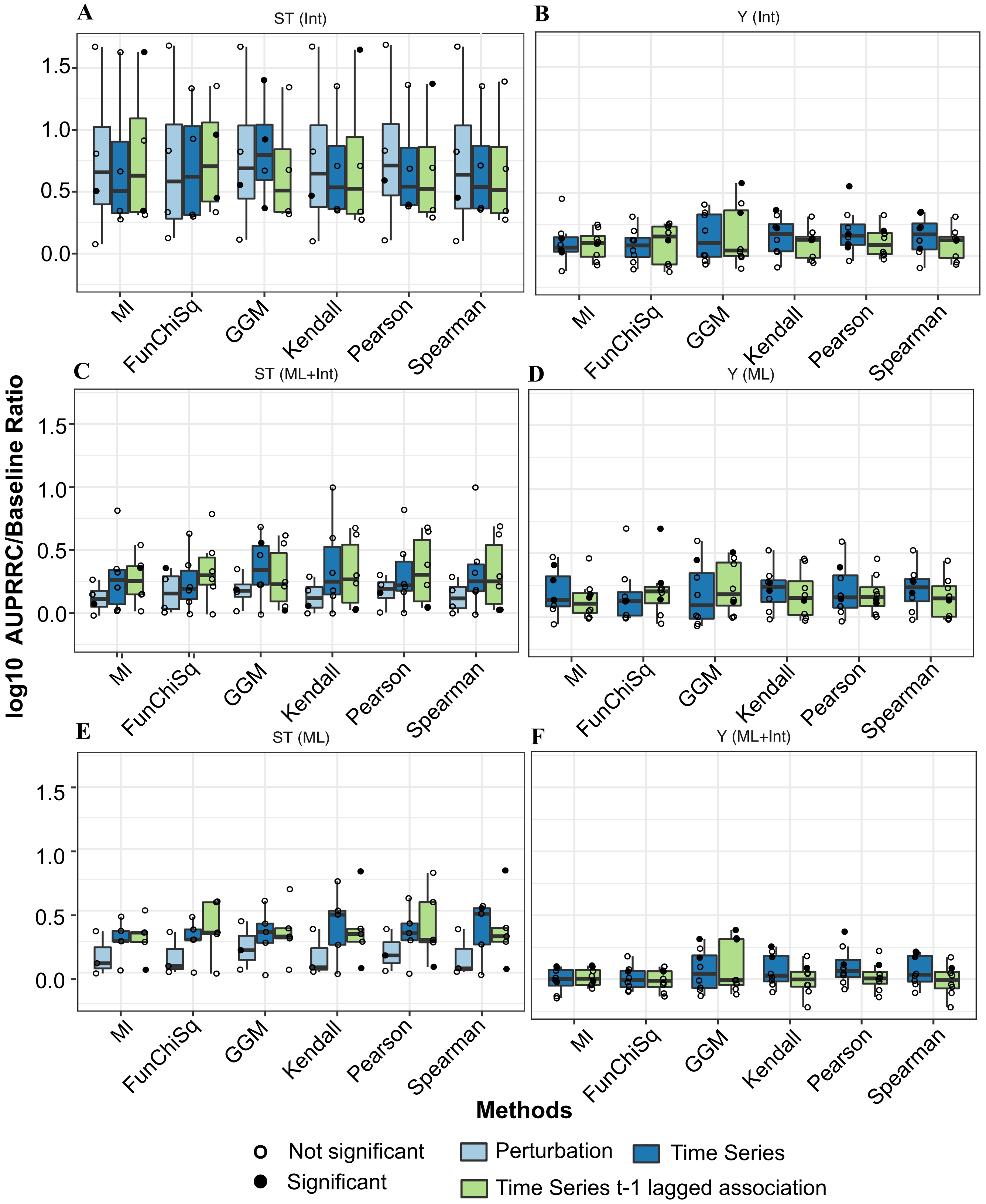


**Figure S3. Distribution plots of Fisher's exact estimated odds ratio for enrichment of true positive hits in top 20% of predicted associations for A.** S/T kinase substrate associations **B.** Y kinase substrate associations **C.** Machine learning derived S/T kinase substrate associations **D.** Machine learning derived Y kinase substrate associations **E.** Combined interactome and machine learning derived S/T kinase substrate associations **F.** Combined interactome and machine learning derivedY kinase substrate associations. The black filled in points on the boxplots indicates statistical significance from the permutation analysis with p-value < 0.05.

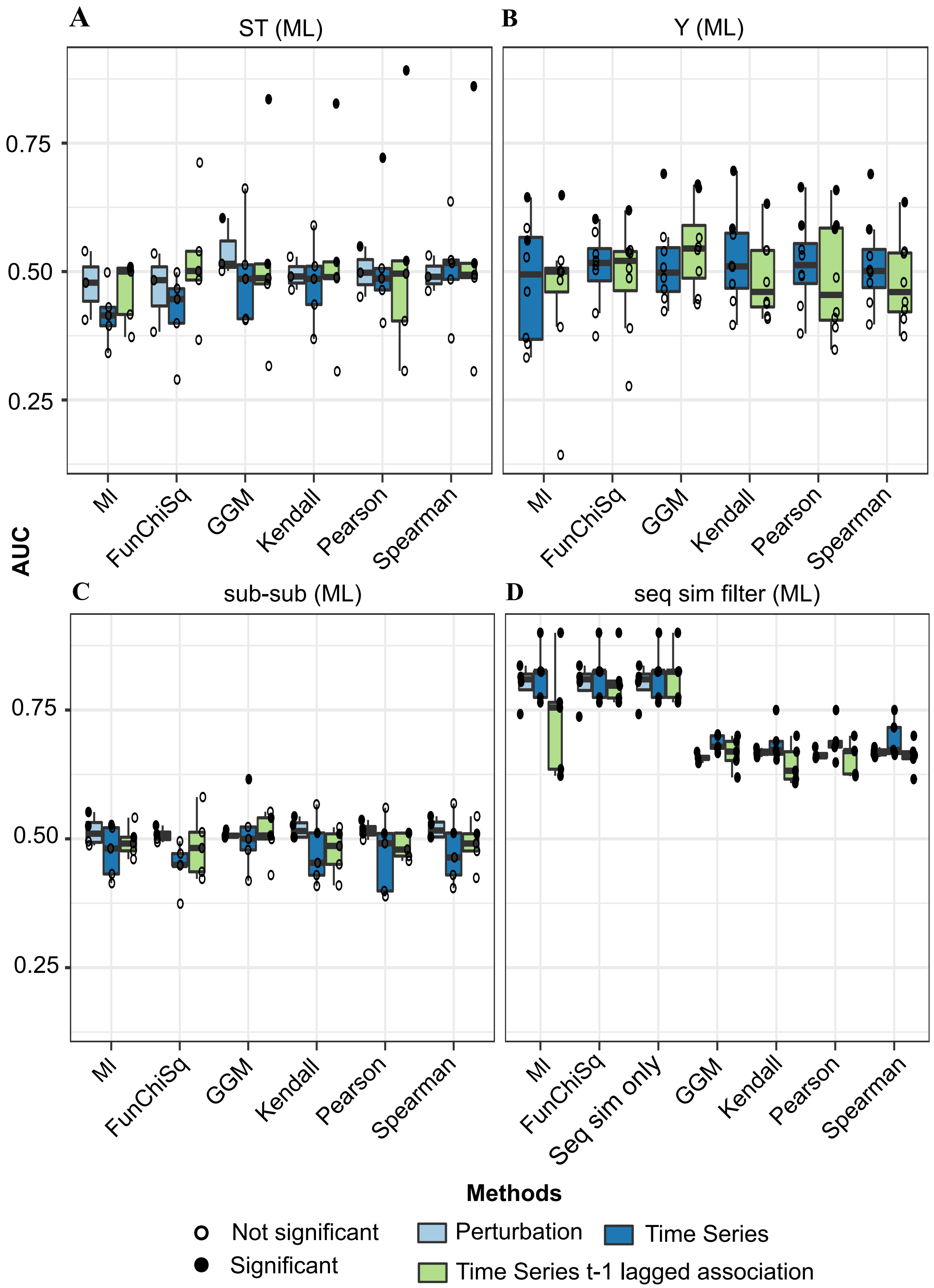


**Figure S4**: **Distribution plots of AUC metrics for A.** S/T kinase substrate associations **B.** Y kinase substrate associations **C.** Pairwise substrate-substrate associations regulated by the same kinase **D.** Pairwise substrate substrate associations regulated by the same kinase with sequence similarity filter. The black filled in points on the boxplots indicates statistical significance from the permutation analysis with p-value < 0.05.

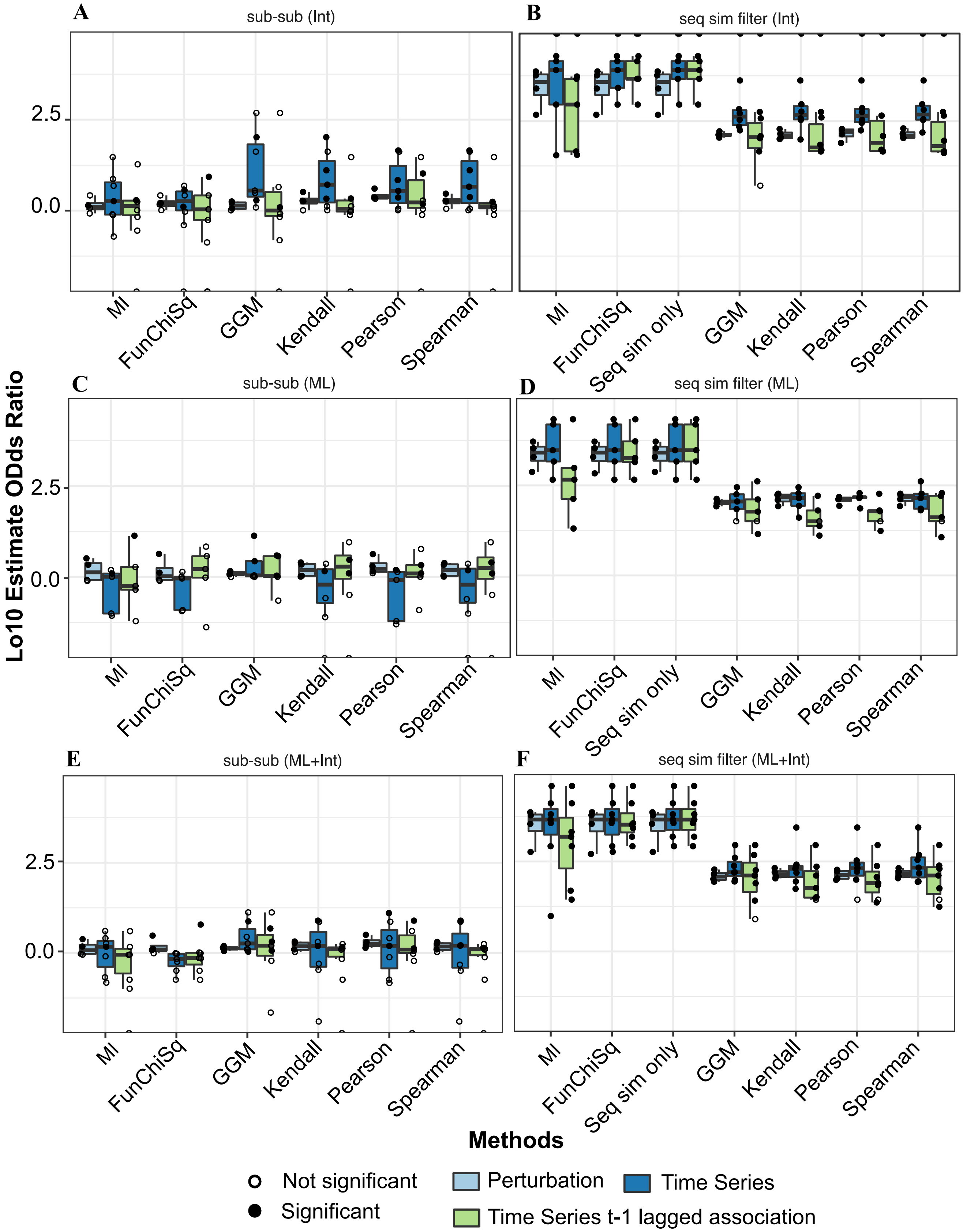


**Figure S5: Distribution plots of log10 estimate odds ratio for interactome derived A.** Substrate-substrate associations **B.** Substrate-substrate associations with sequence similarity filter **C.** Machine learning derived substrate-substrate associations **D.** Machine learning derived substrate-substrate associations with sequence similarity filter **E.** Combined interactome and machine learning derived substrate-substrate associations **F.** Combined interactome and machine learning derived substrate-substrate associations with sequence similarity filter. The black filled in points on the boxplots indicates statistical significance from the permutation analysis with p-value < 0.05.

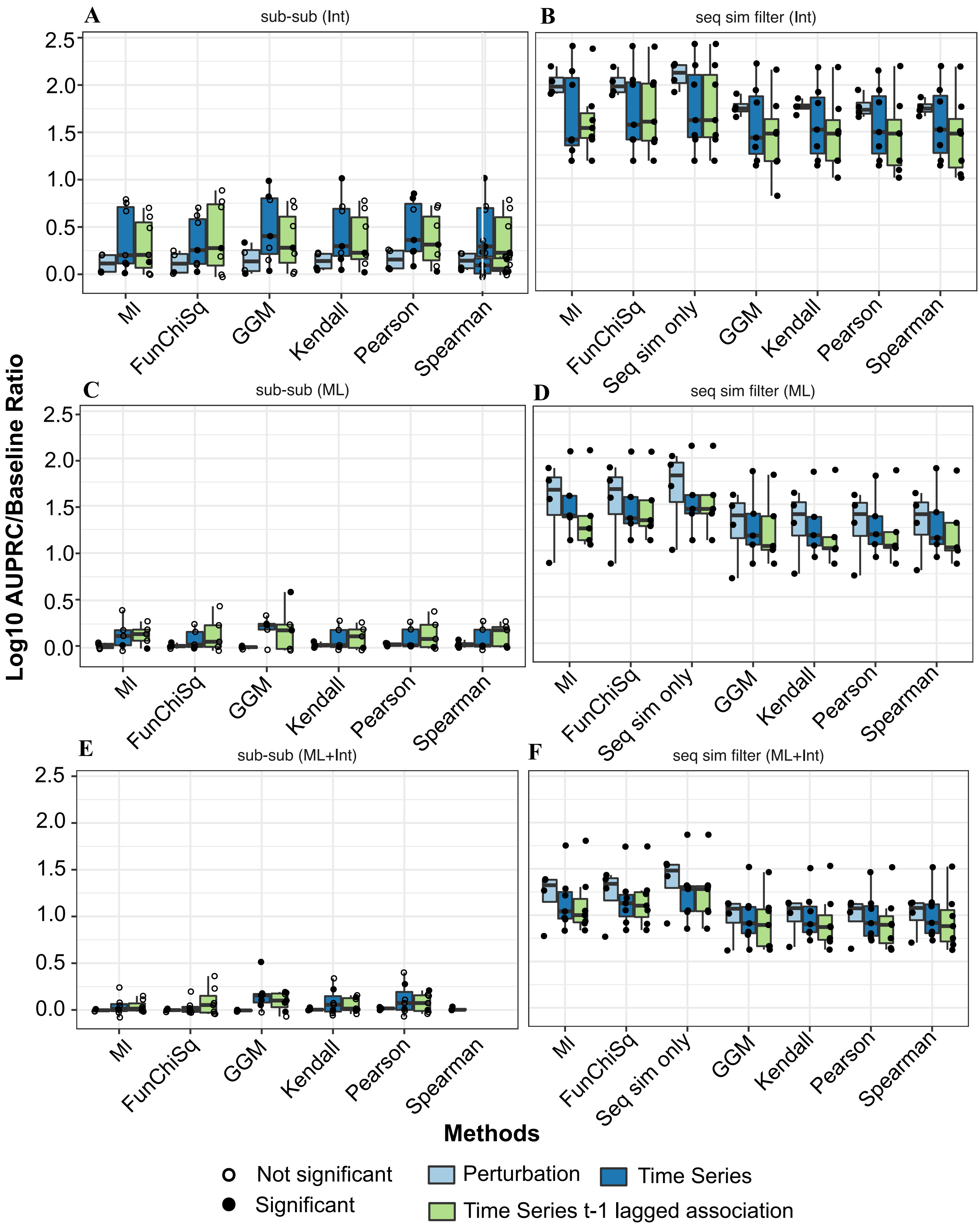


**Figure S6: Distribution plots of log10 AUPRC/baseline ratio for interactome derived A.** Substrate-substrate associations **B.** Substrate-substrate associations with sequence similarity filter **C.** Machine learning derived substrate-substrate associations **D.** Machine learning derived substrate-substrate associations with sequence similarity filter **E.** Combined interactome and machine learning derived substrate-substrate associations **F.** Combined interactome and machine learning derived substrate-substrate associations with sequence similarity filter. The black filled in points on the boxplots indicates statistical significance from the permutation analysis with p-value < 0.05.

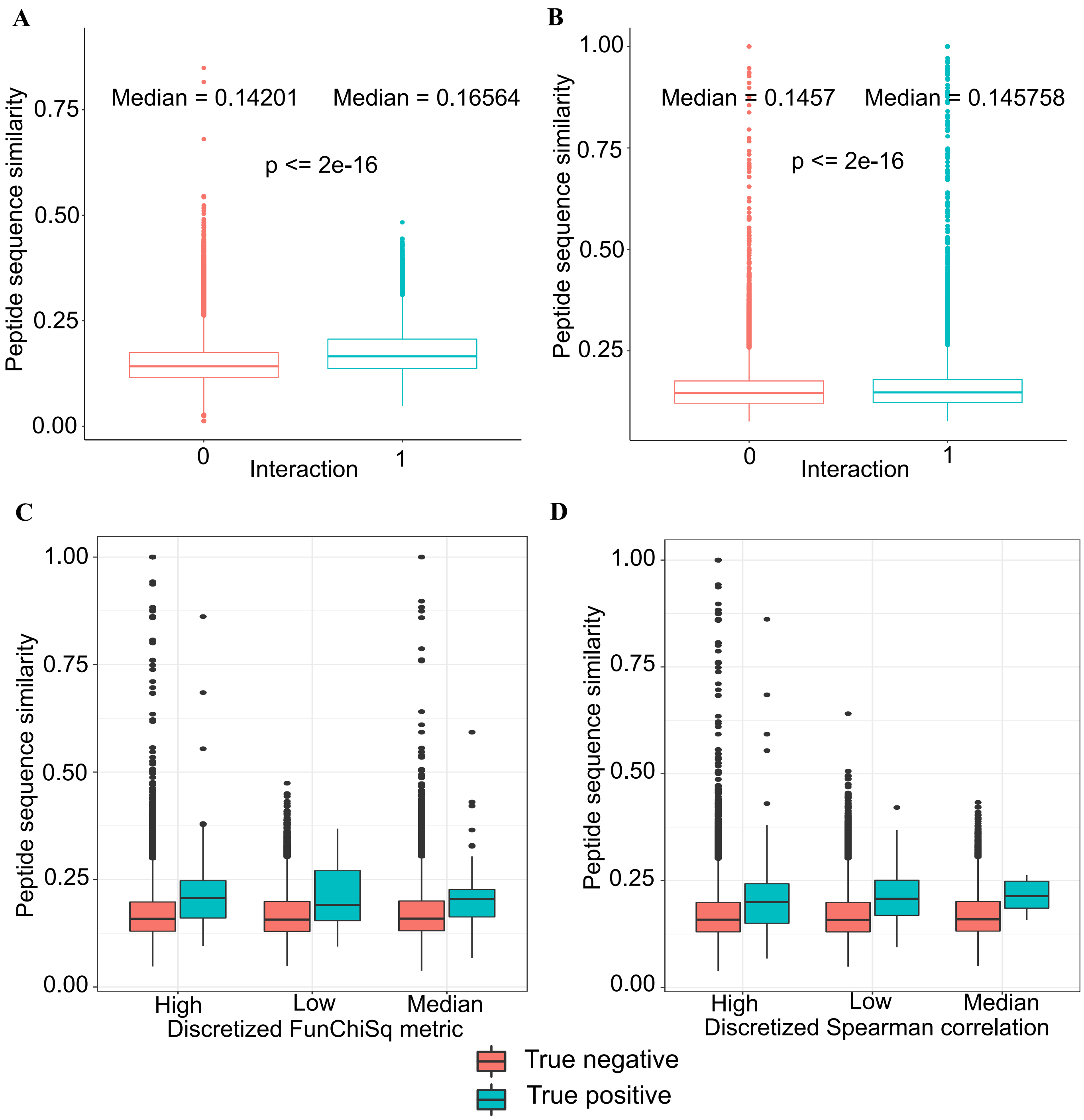


**Figure S7: Distribution of sequence similarity scores between pairs of substrates regulated by the same kinase.** An interaction value of 1 represents a TP, and an interaction value of 0 represents a TN. The pair of distributions plots represent **A.** Serine/threonine phosphosites **B.** Tyrosine phosphosites. Discretized association measures across three levels (High, Medium, Low) with respect to sequence similarity for true positive and true negative interactions from the phosphositeplus database. Plots represent **C.** Discretized FunChiSq **D.** Pearson correlation measures.

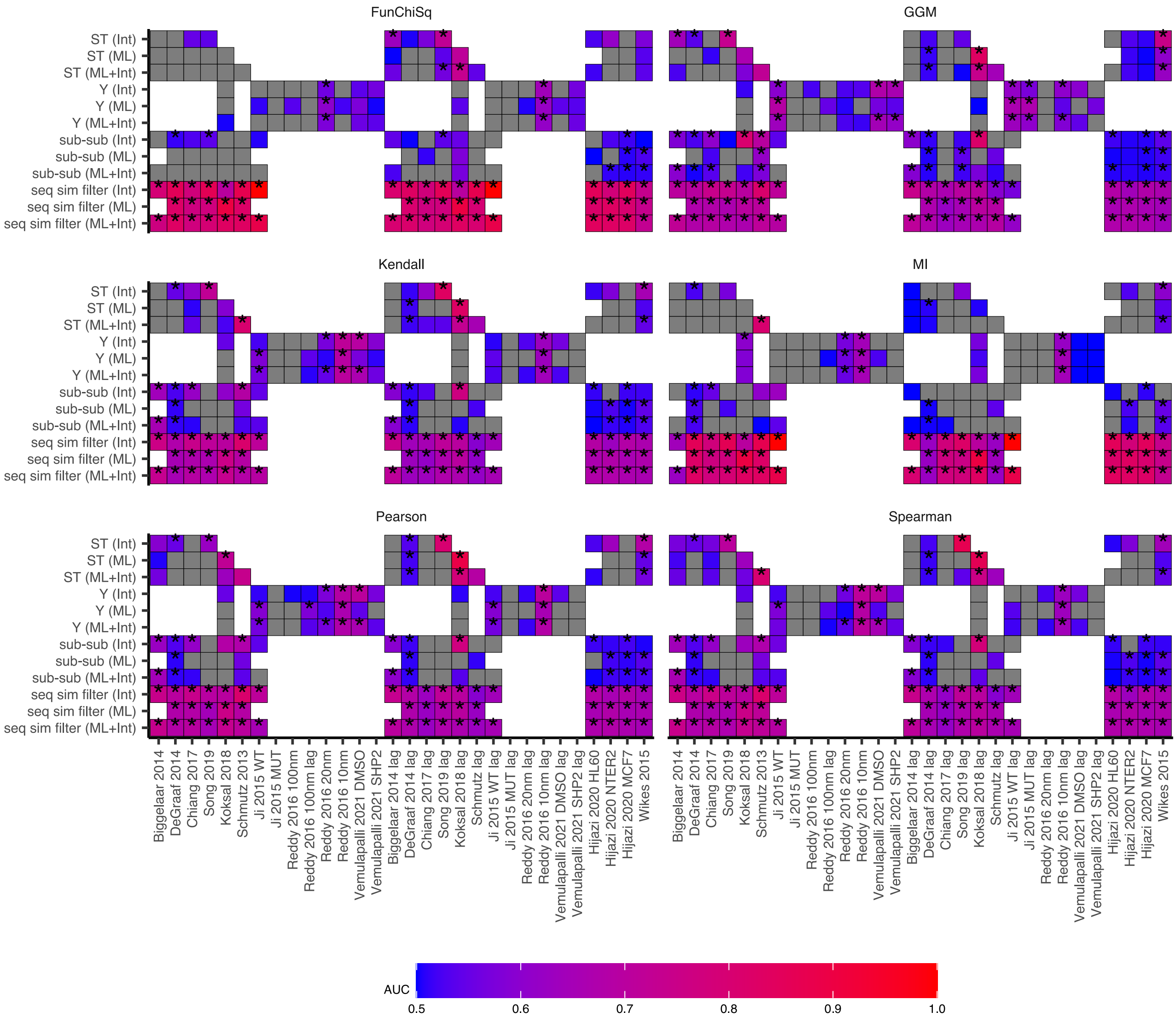


**Figure S8: Heatmap of AUC performance metrics across all datasets and association measrues used for evaluation.** An asterix indicates significance from the permutation analysis with p-value < 0.05. Colourscale of grey indicates datasets with AUC < 0.5.

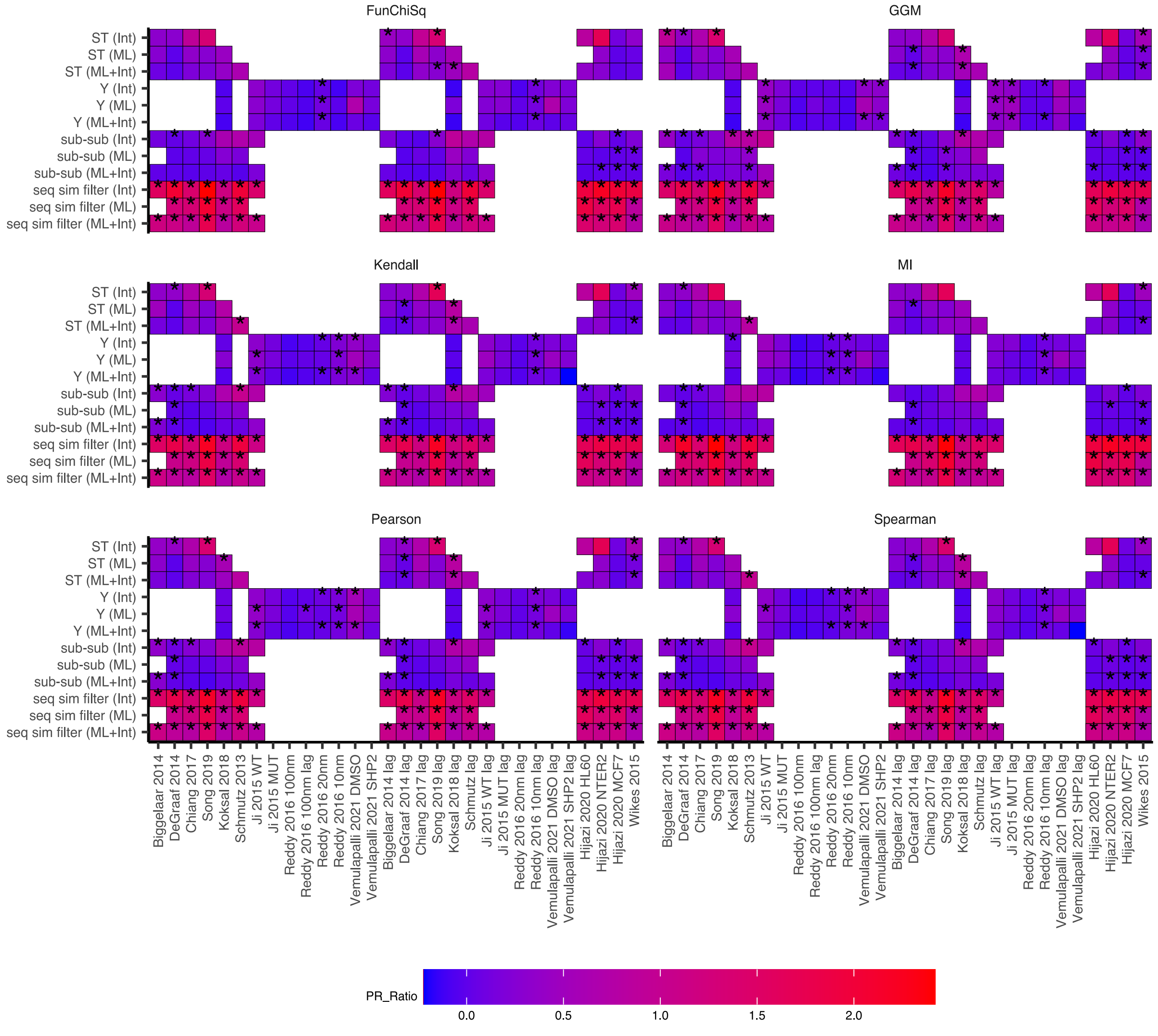


**Figure S9: Heatmap of Precision recall to baseline metric ratio.** The ratio used in the plot has been log10 scaled. An asterix indicates statistical significance from the permutation analysis with p-value < 0.05.

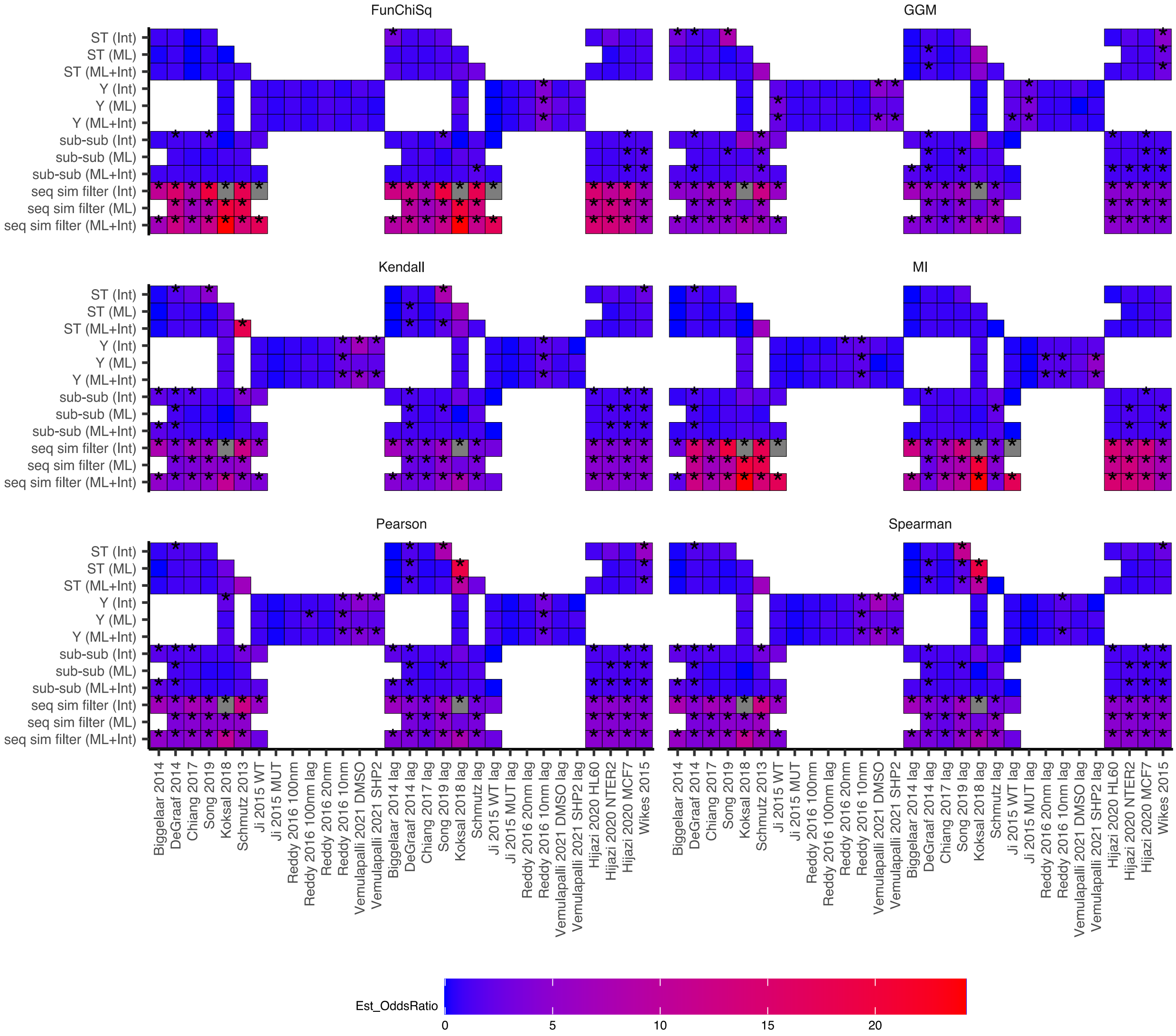


**Figure S10: Heatmap of estimate odds ratio from the Fisher's exact top 20% TP enrichment test.** An asterix indicates statistical significance from the permutation analysis with p-value < 0.05.

**Supplementary Table S1:** Datasets used in this study and the data processing steps applied to each dataset

| Dataset | Reference | Processing |
|---|---|---|
| Koksal et al 2018 | 1 | This dataset was provided as a concatenated dataset across three biological replicates. There are three technical replicates for each biological replicates (3 biological replicates in total). Across the three biological replicates, 1068 peptides were present in all three replicates. This dataset was provided pre-processed (median normalized) as three biological replicates. Each row for each replicate was normalized to the row mean to minimise batch effect variations between the biological replicates. |
| Vemulapalli et al 2021 | 2 | This dataset has 2 replicates, under two conditions: DMSO stimulated, and SHP2 stimulated. These datasets are column median normalised and row mean normalised across each TMT 11plex replicate set. The phosphosites across both replicates were matched. |
| Ji et al 2015 | 3 | Silac labelled cells. Two jurkat cell lines used: mutant (J14–2D1) and WT (J14–76-11). Identical phosphopeptides from the same protein were summed and median normalized again. The data was row mean normalized per replicate set. Replicate 5 was omitted from the analysis as one time point was missing. |
| Song et al 2019 | 4 | This is TMT dataset of a neural progenitor cell line ReN cell line. There are two replicates of the proteome and the phosphoproteoeme. Both datasets were median column nornalised and row mean normalised. A phosphoproteome to proteome ratio for each time point was generated. |
| Biggelaar et al 2014 | 5 | This is a SILAC dataset of thrombin signalling in human endothelial cells. No normalisation or correction is performed on this dataset as each time point sample was provided as a ratio to t=0 stimulation. |

| | | |
|---|---|---|
| De Graaf et al 2014 | 6 | This is a label-free dataset of Jurkat T cells stimulated by prostglandin and measured over six time points. The normalised average signal intensity data was used and a fold change filter was applied to select the differentially expressed phosphosites. Data was normalised to t=0 and analysed as ratios. |
| Reddy et al 2016 | 7 | TMT labelled MCF-10A cells stimulated with different concentrations of EGF (0.2, 0.4, 1, 2.5, 5, 10, 20, 100 nM EGF) measured over 80 seconds. Each dataset had 9 timepoints, but no replicates. The data provided in the supplementary materials was median normalized and normalized to a control channel to generate ratios for each phosphosite. |
| Wilkes et al 2015 | 8 | Label-free datasets. This dataset comes preprocessed with log2 fc ratios against a control sample and p-values. Data was filtered based on the following conditions: 20% or more of the samples in each row needs to have p-value <c0.05 and 3 or more sites are required to have an absolute log2fc ratio greater than 2. |
| Hijazi et al 2020 | 9 | Label-free datasets. This dataset comes preprocessed with log2 fc ratios and p-values. Data filtering steps used for Wilkes et al 2015 was also used for this dataset. |
| Schmutz et al 2013 | 10 | This is a label-free dataset with three replicates in which the phosphoproteome (log2 ratios to t=0) was normalized to the proteome (log2 ratios to t=0) across 4 timepoints. |
| Chiang et al 2017 | 11 | Dataset came as log2 (phos/prot) ratios with three replicates. No processing was required for this dataset. |

### References:

1. Köksal, A. S. *et al.* Synthesizing Signaling Pathways from Temporal Phosphoproteomic Data. *Cell Reports* **24**, 3607–3618 (2018).
2. Vemulapalli, V. *et al.* Time-resolved phosphoproteomics reveals scaffolding and catalysis-responsive patterns of SHP2-dependent signaling. *eLife* **10**, e64251 (2021).
3. Ji, Q., Ding, Y. & Salomon, A. R. SRC Homology 2 Domain-containing Leukocyte Phosphoprotein of 76 kDa (SLP-76) N-terminal Tyrosine Residues Regulate a Dynamic Signaling Equilibrium Involving Feedback of Proximal T-cell Receptor (TCR) Signaling. *Mol Cell Proteomics* **14**, 30–40 (2015).
4. Song, Y. *et al.* A dynamic view of the proteomic landscape during differentiation of ReNcell VM cells, an immortalized human neural progenitor line. *Sci Data* **6**, 190016 (2019).
5. van den Biggelaar, M. *et al.* Quantitative phosphoproteomics unveils temporal dynamics of thrombin signaling in human endothelial cells. *Blood* **123**, e22–e36 (2014).
6. de Graaf, E. L. *et al.* Signal Transduction Reaction Monitoring Deciphers Site-Specific PI3K-mTOR/MAPK Pathway Dynamics in Oncogene-Induced Senescence. *J. Proteome Res.* **14**, 2906–2914 (2015).
7. Reddy, R. J. *et al.* Early signaling dynamics of the epidermal growth factor receptor. *Proc Natl Acad Sci U S A* **113**, 3114–3119 (2016).
8. Wilkes, E. H., Terfve, C., Gribben, J. G., Saez-Rodriguez, J. & Cutillas, P. R. Empirical inference of circuitry and plasticity in a kinase signaling network. *PNAS* **112**, 7719–7724 (2015).
9. Hijazi, M., Smith, R., Rajeeve, V., Bessant, C. & Cutillas, P. R. Reconstructing kinase network topologies from phosphoproteomics

data reveals cancer-associated rewiring. *Nature Biotechnology* **38**, 493–502 (2020).

10. Schmutz, C. *et al.* Systems-Level Overview of Host Protein Phosphorylation During Shigella flexneri Infection Revealed by Phosphoproteomics. *Molecular & Cellular Proteomics* **12**, 2952–2968 (2013).

11. Chiang, C.-K. *et al.* Quantitative phosphoproteomics reveals involvement of multiple signaling pathways in early phagocytosis by the retinal pigmented epithelium. *Journal of Biological Chemistry* **292**, 19826–19839 (2017).
